# Effect of SARS-CoV-2 proteins on vascular permeability

**DOI:** 10.1101/2021.02.27.433186

**Authors:** Rossana Rauti, Meishar Shahoha, Yael Leichtmann-Bardoogo, Rami Nasser, Rina Tamir, Victoria Miller, Tal Babich, Kfir Shaked, Avner Ehrlich, Konstantinos Ioannidis, Yaakov Nahmias, Roded Sharan, Uri Ashery, Ben M. Maoz

**Author notes:** These authors are equal contributors (Rossana Rauti, Meishar Shahoha, Yael Leichtmann-Bardoogo). Corresponding authors Uri Ashery, mail, Ben M. Maoz, mail.

## Abstract

SARS-CoV-2 infection leads to severe disease associated with cytokine storm, vascular dysfunction, coagulation, and progressive lung damage. It affects several vital organs, seemingly through a pathological effect on endothelial cells. The SARS-CoV-2 genome encodes 29 proteins, whose contribution to the disease manifestations, and especially endothelial complications, is unknown. We cloned and expressed 26 of these proteins in human cells and characterized the endothelial response to overexpression of each, individually. Whereas most proteins induced significant changes in endothelial permeability, nsp2, nsp5_c145a (catalytic dead mutant of nsp5) and nsp7 also reduced CD31, and increased von Willebrand factor expression and IL-6, suggesting endothelial dysfunction. Using propagation-based analysis of a protein–protein interaction (PPI) network, we predicted the endothelial proteins affected by the viral proteins that potentially mediate these effects. We further applied our PPI model to identify the role of each SARS-CoV-2 protein in other tissues affected by COVID-19. Overall, this work identifies the SARS-CoV-2 proteins that might be most detrimental in terms of endothelial dysfunction, thereby shedding light on vascular aspects of COVID-19.

## Introduction

Coronavirus disease (COVID-19) caused by the 2019 novel coronavirus (2019-nCoV/SARS-CoV-2) led to a global pandemic in 2020. By early February 2021, coronavirus had infected more than 105 million people worldwide, causing over 2.3 million deaths. After the initial phase of the viral infection, ~30% of patients hospitalized with COVID-19 develop severe disease with progressive lung damage, known as severe acute respiratory syndrome (SARS), and a severe immune response. Interestingly, additional pathologies have been observed, such as hypoxemia and cytokine storm which, in some cases, lead to heart and kidney failure, and neurological symptoms. Recent observations suggest that these pathologies are mainly due to increased coagulation and vascular dysfunction^1–3^. It is currently believed that in addition to being a respiratory disease, COVID-19 might also be a “vascular disease”^1^, as it may result in a leaky vascular barrier and increased expression of von Willebrand factor (VWF)3, responsible for increased coagulation, cytokine release, and inflammation^3–12^. Recent studies suggest that the main mechanism disrupting the endothelial barrier occurs in several stages: **(a)** a direct effect on the endothelial cells that causes endotheliitis and endothelial dysfunction, **(b)** lysis and death of the endothelial cells^4,12^, **(c)** sequestering of human angiotensin I-converting enzyme 2 (hACE2) by viral spike proteins that activates the kallikrein–bradykinin and renin–angiotensin pathways, increasing vascular permeability^4,13^, and **(d)** overreaction of the immune system, during which a combination of neutrophils and immune cells producing reactive oxygen species, inflammatory cytokines (e.g., interleukin [IL]-1β, IL-6 and tumor necrosis factor) and vasoactive molecules (e.g., thrombin, histamine, thromboxane A2 and vascular endothelial growth factor), and the deposition of hyaluronic acid lead to disruption of endothelial junctions, increased vascular permeability, and leakage and coagulation^2,4,13^. Of great interest is the effect on the brain’s vascular system. Cerebrovascular effects have been suggested to be among the long-lasting effects of COVID-19. Indeed, the susceptibility of brain endothelial cells to direct SARS-CoV-2 infection was found to increase due to increased expression of hACE2 in a flow-dependent manner, leading to a unique gene-expression process that might contribute to the cerebrovascular effects of the virus^14^.

While many studies point out the importance of the vascular system in COVID-19^15–17^, only a few^18–21^ have looked at the direct vascular response to the virus. Most of those reports stem from either clinical observations, or *in-vitro* or *in-vivo* studies in which animals/cells were transfected with the SARS-CoV-2 virus and their systemic cellular response assessed, without pinpointing the specific viral protein(s) causing the observed changes. SARS-CoV-2 is an enveloped virus with a positive-sense, single-stranded RNA genome of ~30 kb, which encodes 29 proteins (**Fig. 1**). These proteins can be classified as: *structural proteins*: S (spike proteins), E (envelope proteins), M (membrane proteins), N (nucleocapsid protein and viral RNA); *nonstructural proteins*: nsp1–16; *open reading frame accessory proteins*: orf3–10^22,23^. **Table 1** summarizes the known effects of specific SARS-CoV-2 proteins^24^. The functionality of some of these is still not known. Moreover, there remains a large lack of knowledge on the molecular mechanisms, especially the protein–protein interaction (PPI) pathways^25^, leading to tissue dysfunction.

**Fig. 1.**
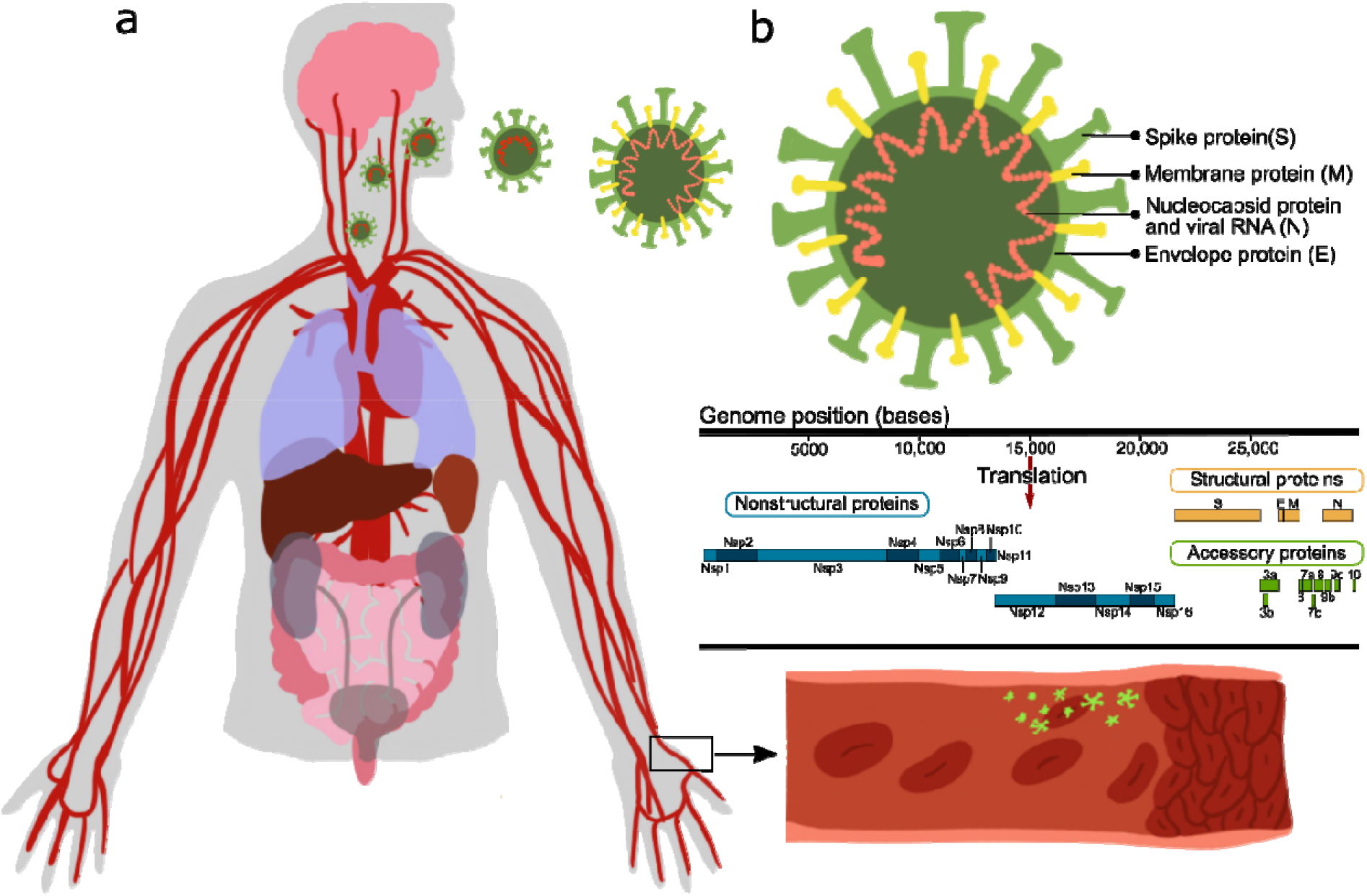
Effect of SARS-CoV-2 proteins on endothelial cells. **a** Sketch representing the main organs affected by SARS-CoV-2; **b** structure and gene composition of SARS-CoV-2.

**Table 1.**
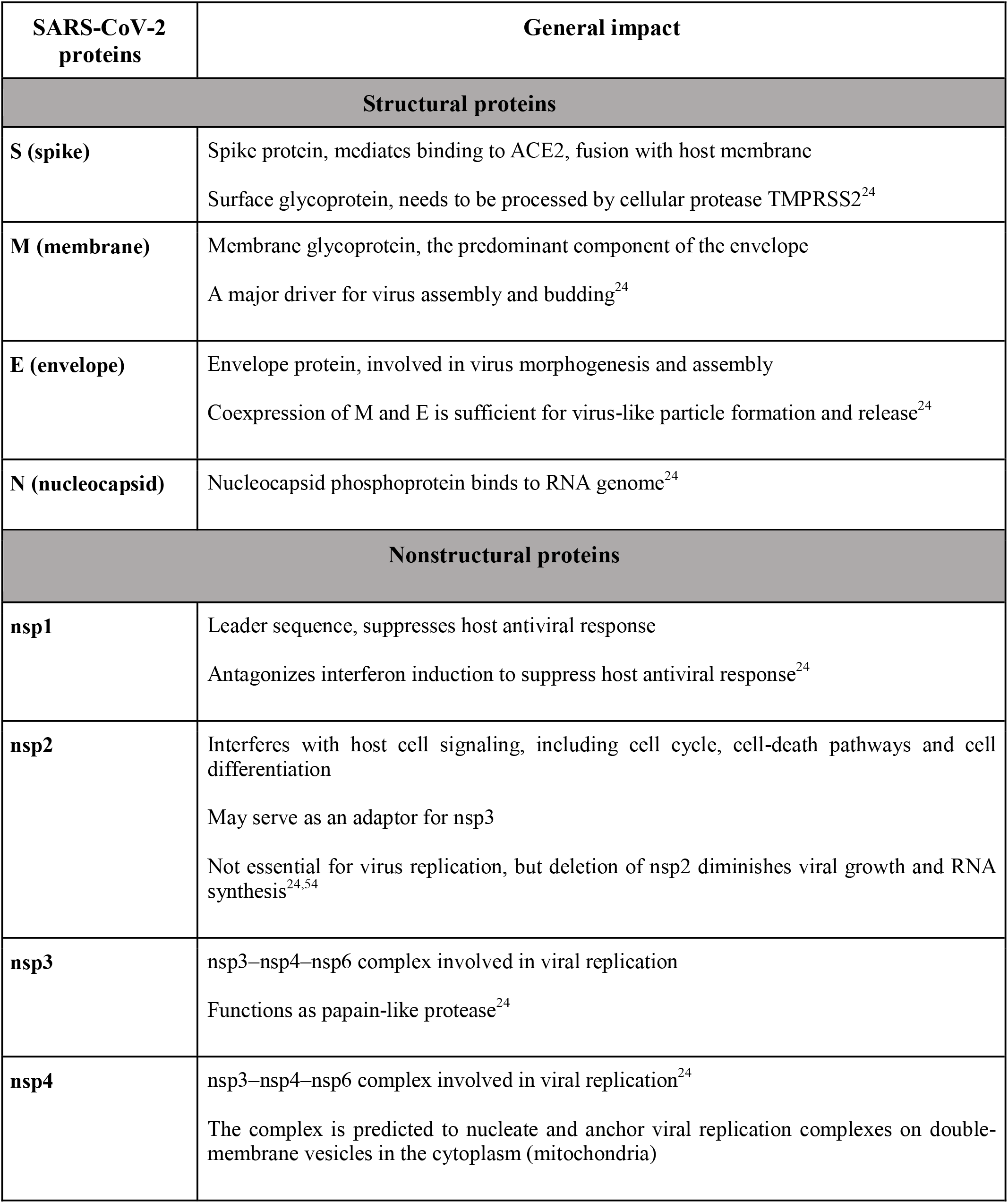
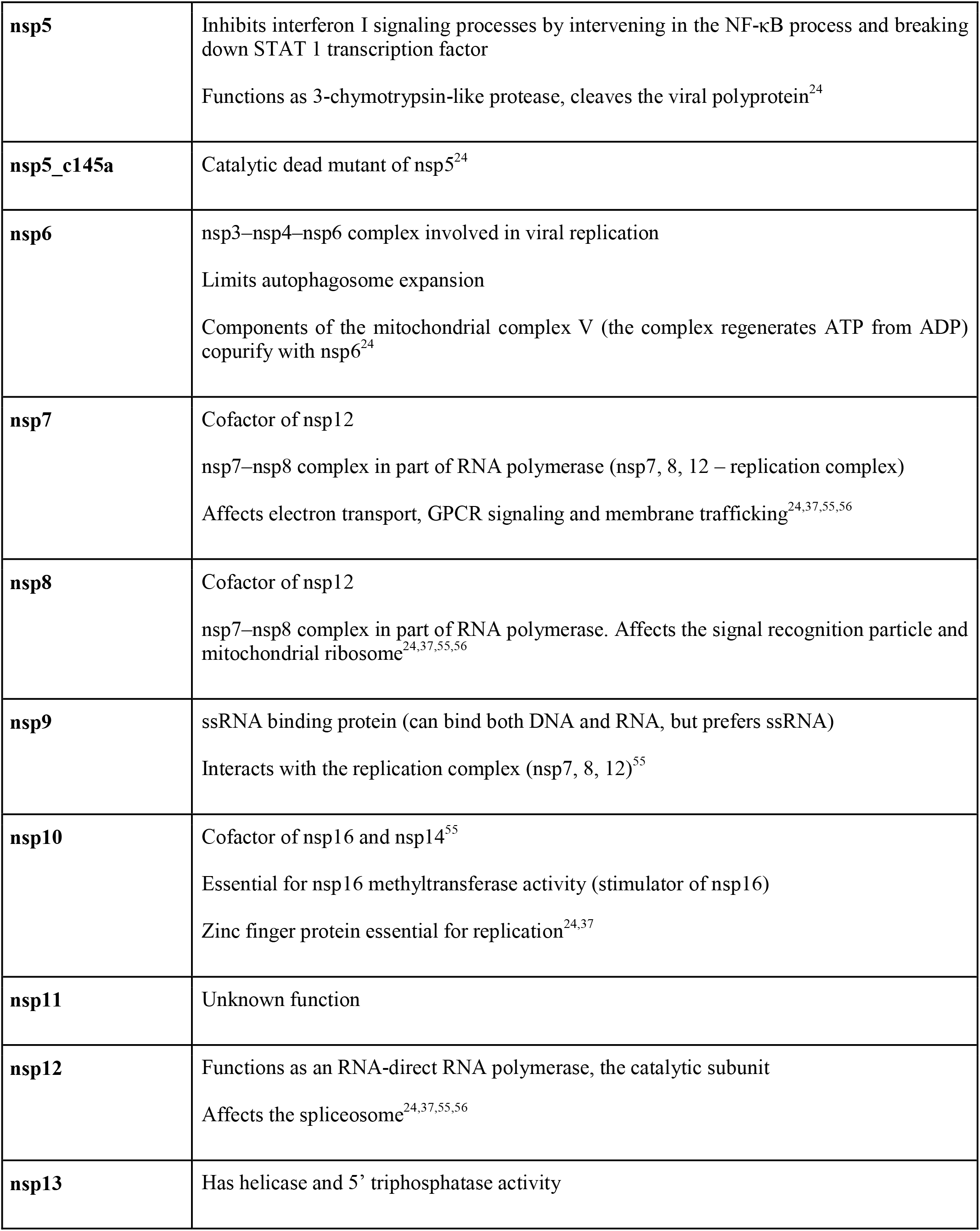
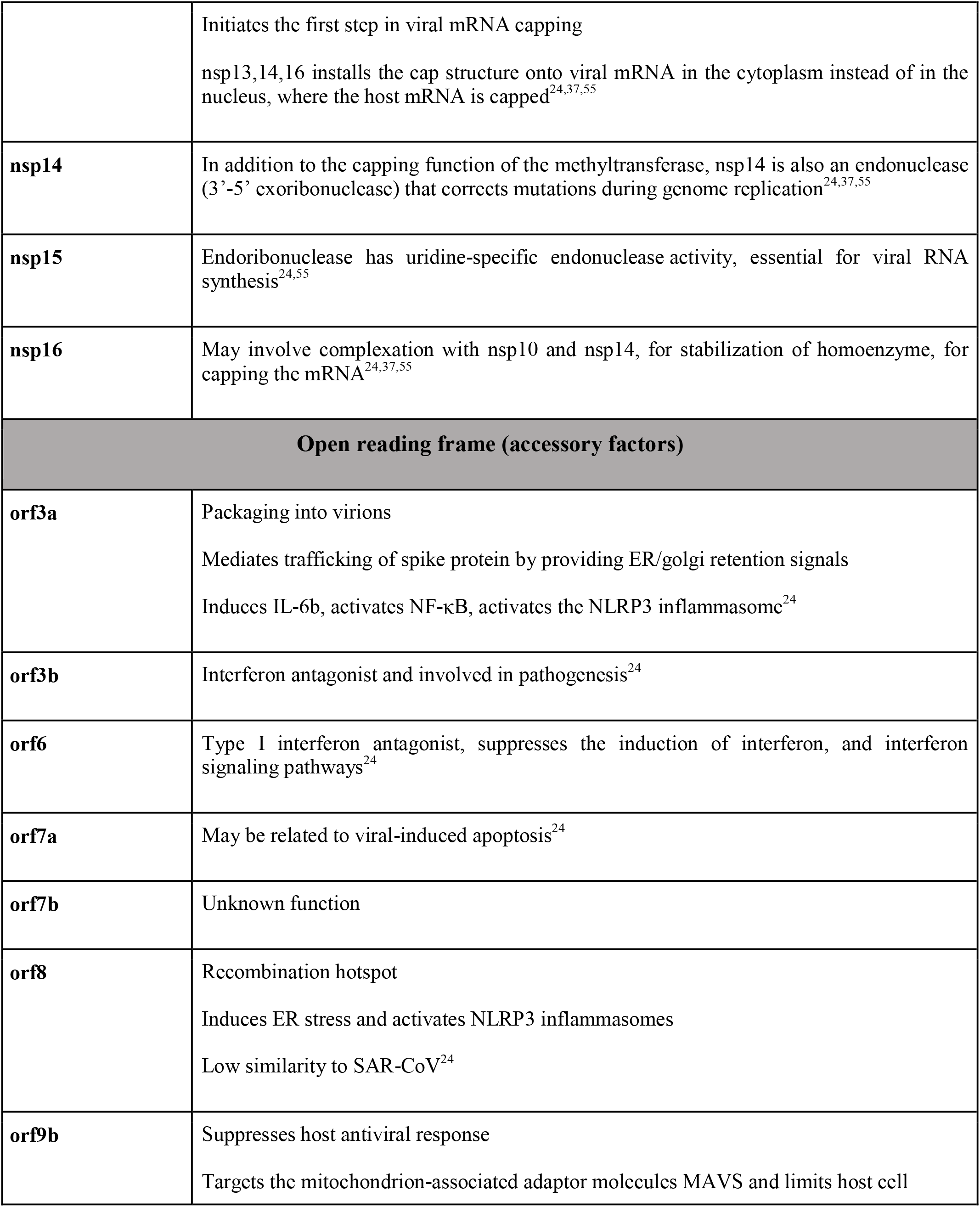
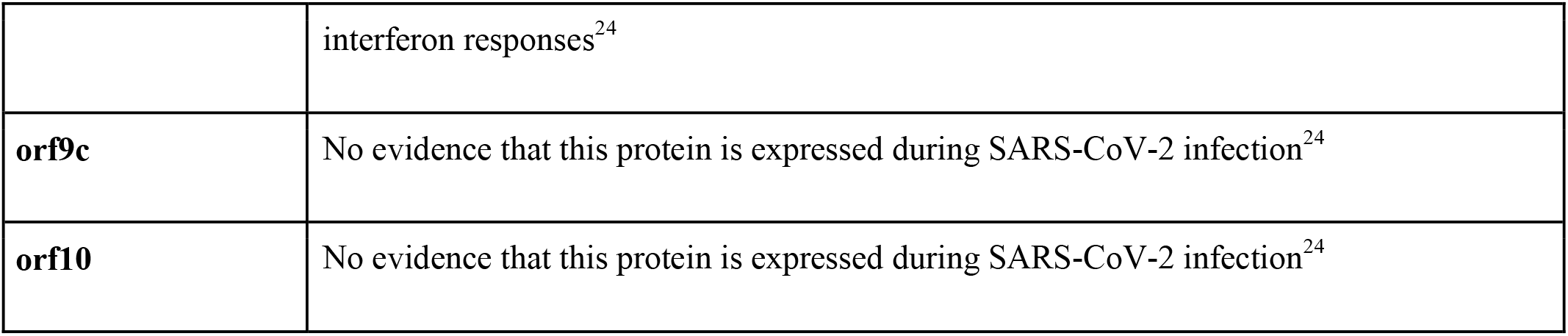
SARS-CoV-2 proteins.

| **SARS-CoV-2 proteins** | **General impact** |
| --- | --- |
| **Structural proteins** |  |
| **S (spike)** | Spike protein, mediates binding to ACE2, fusion with host membrane<br>Surface glycoprotein, needs to be processed by cellular protease TMPRSS2<sup>24</sup> |
| **M (membrane)** | Membrane glycoprotein, the predominant component of the envelope<br>A major driver for virus assembly and budding<sup>24</sup> |
| **E (envelope)** | Envelope protein, involved in virus morphogenesis and assembly<br>Coexpression of M and E is sufficient for virus-like particle formation and release<sup>24</sup> |
| **N (nucleocapsid)** | Nucleocapsid phosphoprotein binds to RNA genome<sup>24</sup> |
| **Nonstructural proteins** |  |
| **nsp1** | Leader sequence, suppresses host antiviral response<br>Antagonizes interferon induction to suppress host antiviral response<sup>24</sup> |
| **nsp2** | Interferes with host cell signaling, including cell cycle, cell-death pathways and cell differentiation<br>May serve as an adaptor for nsp3<br>Not essential for virus replication, but deletion of nsp2 diminishes viral growth and RNA synthesis<sup>24,54</sup> |
| **nsp3** | nsp3–nsp4–nsp6 complex involved in viral replication<br>Functions as papain-like protease<sup>24</sup> |
| **nsp4** | nsp3–nsp4–nsp6 complex involved in viral replication<sup>24</sup><br>The complex is predicted to nucleate and anchor viral replication complexes on double-membrane vesicles in the cytoplasm (mitochondria) |
| **nsp5** | Inhibits interferon I signaling processes by intervening in the NF-κB process and breaking down STAT 1 transcription factor<br>Functions as 3-chymotrypsin-like protease, cleaves the viral polyprotein<sup>24</sup> |
| **nsp5_c145a** | Catalytic dead mutant of nsp5<sup>24</sup> |
| **nsp6** | nsp3–nsp4–nsp6 complex involved in viral replication<br>Limits autophagosome expansion<br>Components of the mitochondrial complex V (the complex regenerates ATP from ADP) copurify with nsp6<sup>24</sup> |
| **nsp7** | Cofactor of nsp12<br>nsp7–nsp8 complex in part of RNA polymerase (nsp7, 8, 12 – replication complex)<br>Affects electron transport, GPCR signaling and membrane trafficking<sup>24,37,55,56</sup> |
| **nsp8** | Cofactor of nsp12<br>nsp7–nsp8 complex in part of RNA polymerase. Affects the signal recognition particle and mitochondrial ribosome<sup>24,37,55,56</sup> |
| **nsp9** | ssRNA binding protein (can bind both DNA and RNA, but prefers ssRNA)<br>Interacts with the replication complex (nsp7, 8, 12)<sup>55</sup> |
| **nsp10** | Cofactor of nsp16 and nsp14<sup>55</sup><br>Essential for nsp16 methyltransferase activity (stimulator of nsp16)<br>Zinc finger protein essential for replication<sup>24,37</sup> |
| **nsp11** | Unknown function |
| **nsp12** | Functions as an RNA-direct RNA polymerase, the catalytic subunit<br>Affects the spliceosome<sup>24,37,55,56</sup> |
| **nsp13** | Has helicase and 5' triphosphatase activity |
|  | Initiates the first step in viral mRNA capping<br>nsp13,14,16 installs the cap structure onto viral mRNA in the cytoplasm instead of in the nucleus, where the host mRNA is capped<sup>24,37,55</sup> |
| **nsp14** | In addition to the capping function of the methyltransferase, nsp14 is also an endonuclease (3'-5' exoribonuclease) that corrects mutations during genome replication<sup>24,37,55</sup> |
| **nsp15** | Endoribonuclease has uridine-specific endonuclease activity, essential for viral RNA synthesis<sup>24,55</sup> |
| **nsp16** | May involve complexation with nsp10 and nsp14, for stabilization of homoenzyme, for capping the mRNA<sup>24,37,55</sup> |
| **Open reading frame (accessory factors)** |  |
| **orf3a** | Packaging into virions<br>Mediates trafficking of spike protein by providing ER/golgi retention signals<br>Induces IL-6b, activates NF-κB, activates the NLRP3 inflammasome<sup>24</sup> |
| **orf3b** | Interferon antagonist and involved in pathogenesis<sup>24</sup> |
| **orf6** | Type I interferon antagonist, suppresses the induction of interferon, and interferon signaling pathways<sup>24</sup> |
| **orf7a** | May be related to viral-induced apoptosis<sup>24</sup> |
| **orf7b** | Unknown function |
| **orf8** | Recombination hotspot<br>Induces ER stress and activates NLRP3 inflammasomes<br>Low similarity to SAR-CoV<sup>24</sup> |
| **orf9b** | Suppresses host antiviral response<br>Targets the mitochondrion-associated adaptor molecules MAVS and limits host cell |
|  | interferon responses<sup>24</sup> |
| **orf9c** | No evidence that this protein is expressed during SARS-CoV-2 infection<sup>24</sup> |
| **orf10** | No evidence that this protein is expressed during SARS-CoV-2 infection<sup>24</sup> |

To tackle these challenges, we cultured human umbilical vein endothelial cells (HUVEC) and transduced them with lentiviral particles encoding each of 26 of the viral proteins, separately. We then examined their effects on HUVEC monolayer permeability and the expression of factors involved in vascular permeability and coagulation. The results were analyzed in the context of virus–host and host–host PPI networks. By combining the insights from the experimental and computational results, we generated a model that explains how each of the 26 proteins of SARS-CoV-2, including a mutated form of nsp5, the catalytic dead mutant termed nsp5_c145a, affects the protein network regulating vascular functionality. Moreover, once the PPI model was validated with our experimental data, we applied it to more than 250 proteins that have been identified in the literature as affected by the SARS-CoV-2 proteins. This enabled us to pinpoint the more dominant SARS-CoV-2 proteins and chart their effects. Overall, this work shows how each of the SARS-CoV-2 proteins affects vascular functionality; moreover, once the model was validated, we applied it to identify how SARS-CoV-2 proteins interact with proteins which have been significantly correlated with changes in cell functionality.

## Results

Increasing numbers of studies indicate a significant role for the vasculature in the physiological response to SARS-CoV-2. However, neither the exact molecular mechanism that leads to these effects nor the individual contribution of any of the SARS-CoV-2 proteins is known. Plasmids encoding SARS-CoV-2 proteins were cloned into lentivirus vectors, with eGFP-encoding vector used as a negative control (*Methods*). To shed light on the vascular response to the virus, HUVEC were cultured on different platforms, transduced with these lentiviral particles, and assessed for the effects of the virus proteins on different functionalities. Culturing HUVEC on Transwells (**Fig. 2a**) allowed us to identify how the specific proteins affect endothelial functionality. To ensure proper transfection, the control vector included a GFP label, which enabled us to estimate transfection efficiency at around 70% (**Fig. 2a**). Since the most basic function of the endothelium is to serve as a barrier, we sought to identify the changes in endothelium permeability in response to the SARS-CoV-2 proteins, and to pinpoint which of these proteins have the most significant effect. Permeability was measured via trans-epithelial-endothelial electrical resistance (TEER), a standard method that identifies changes in impedance values. The GFP control and 9 SARS-CoV-2 proteins did not show any significant change in TEER values (compared to the untreated condition), whereas 18 of the SARS-CoV-2 proteins caused significant changes in value (see plot in **Fig. 2b**). The most dominant permeability changes were observed with nsp5_c145a, nsp13, nsp7, orf7a and nsp2, with a 20–28% decrease in TEER values (**Fig. 2b**, and **Fig. 2c**, in which the different SARS-CoV-2 proteins are listed and the gradual color change from red to violet represents the progressive reduction in TEER values). Next, we analyzed some of the proteins that exhibited the most significant (nsp2, nsp5_c145a and nsp7) or least significant (S) changes in TEER value for changes in expression of the cell-junction protein CD31, indicating altered permeability (**Fig. 2d,e)**. Quantification of the immunohistochemistry (IHC) (**Fig. 2d,e)** showed, as expected, that nsp2, nsp5_c145a and nsp7 significantly reduce the expression levels of CD31 compared to the untreated, eGFP and S conditions, suggesting a deterioration in barrier function. Hence, these data show a differential effect of SARS-CoV-2 proteins on endothelial functionality and provide a mechanistic explanation for the reduction in endothelial integrity.

**Fig. 2.**
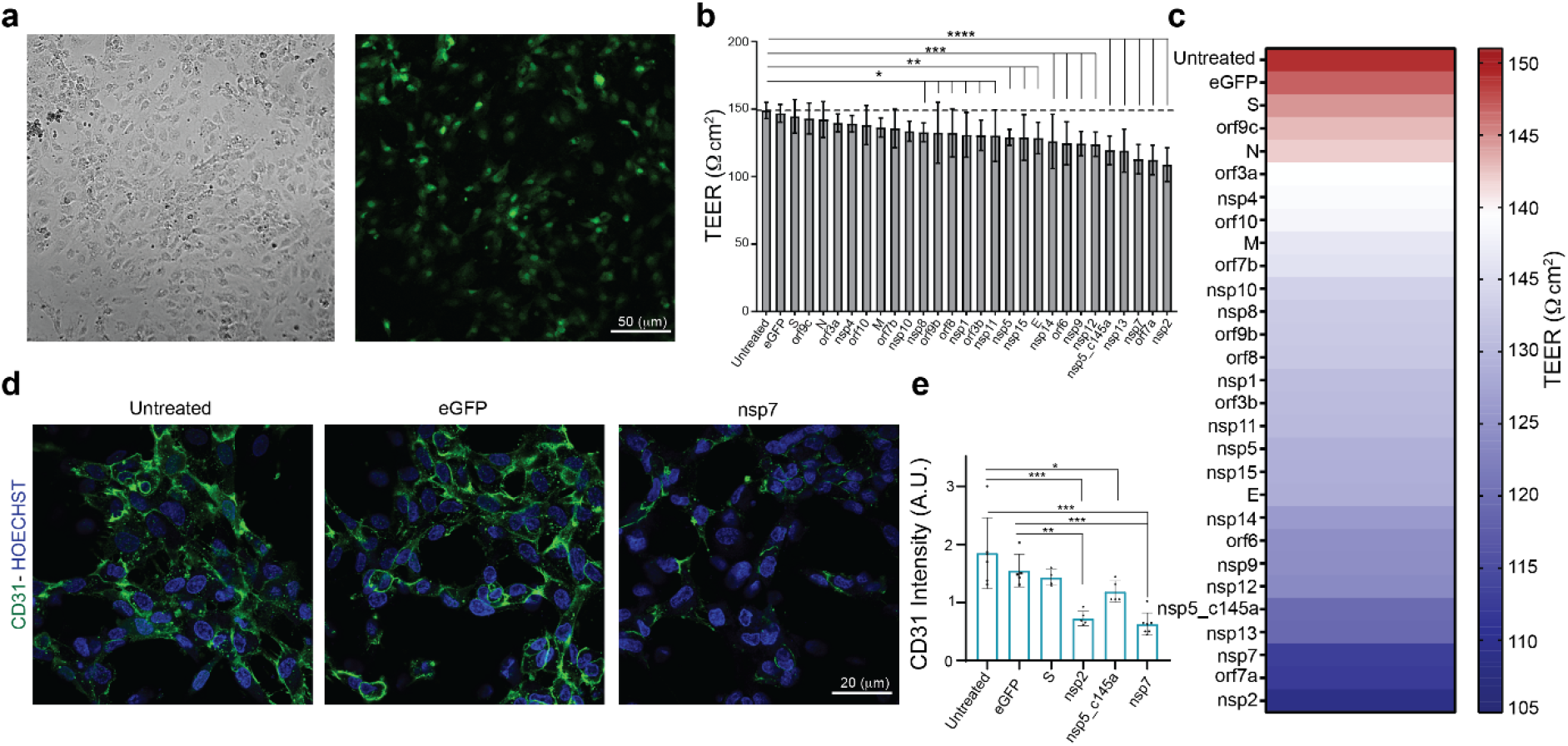
Effect of SARS-CoV-2 proteins on HUVEC. **a** Bright-field and fluorescent image of infected eGFP HUVEC, scale bar: 50 μm; **b** permeability changes as a result of SARS-CoV-2 proteins were assessed by TEER measurement. Note the statistical differences compared to the untreated control condition; **c** color map showing a gradual decrease in TEER values compared to the untreated condition; **d** IHC for CD31 (green) and Hoechst (blue) for the three specified conditions, scale bar: 20 μm; **e** quantification of CD31 expression levels.

It is now known that SARS-CoV-2 causes a severe cytokine storm^8,26^ and a significant increase in coagulation-related pathologies. As we were interested in identifying the role of the vasculature in these observations, we stained and quantified the expression level of VWF (**Fig. 3a,b**), which is highly correlated with coagulation^27^. Similar to the CD31 staining, we characterized only those proteins that resulted in a significant decrease in TEER values (nsp2, nsp5_c145a and nsp7). As shown in **Fig. 3a,b**, the control samples did not exhibit marked expression of VWF, whereas the cells transfected with nsp2, nsp5_c145a and nsp7 showed a significant change in VWF expression. Moreover, as VWF is also associated with increased inflammation^28^, we monitored changes in cytokine expression due to the different SARS-CoV-2 proteins (**Fig. 3c)**. We were particularly interested in IL-6, which has been identified as one of the most dominant cytokines expressed as a result of SARS-CoV-2 infection^26,29–32^. We observed that 13 out of the 26 proteins caused an increase in IL-6 secretion, 3 of which had resulted in a decrease in barrier function and increased VWF expression.

**Fig. 3.**
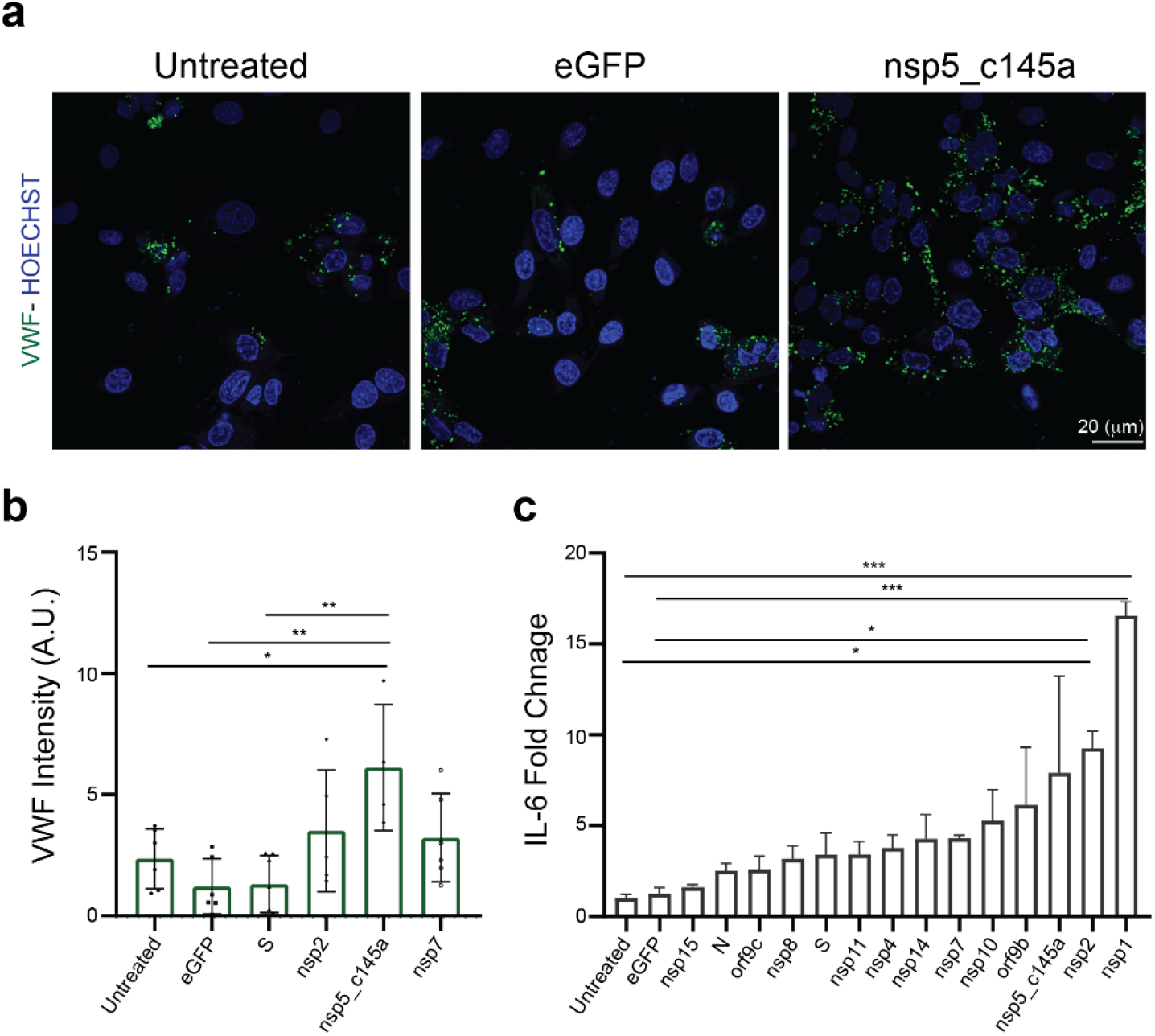
HUVEC response to specific proteins. **a** Confocal reconstructions of HUVEC stained for VWF (green) and Hoechst (blue) for three conditions: control (untreated), eGFP, and nsp5_c145a, scale bar: 20 μm; **b** quantification of VWF expression levels; **c** fold change of IL-6 in response to the different proteins.

We then investigated how SARS-CoV-2 causes the observed changes in HUVEC permeability. We collected sets of proteins responsible for specific functionalities of endothelial cells. We also constructed an integrated viral–host and host–host PPI network. For each viral protein and each prior functional set, we measured the network proximity between the viral protein and the human functional set using a network propagation algorithm. We scored the significance of these propagation calculations by comparing them to those obtained on random PPI networks with the same node degrees. Proteins receiving high and significant scores were most likely to interact with the specific SARS-CoV-2 protein and thus might cause the observed functional changes. When comparing the overall effects of the 26 SARS-CoV-2 proteins on endothelial tight-junction proteins (e.g., cadherin 1–5, occludin and ZO 1–3), we found a correlation between the effects of the SARS-CoV-2 proteins and TEER values (**Fig. 4a**). Moreover, some of the proteins that significantly affected the TEER parameters (**Fig. 2c)** were also observed to be significantly proximal to the permeability-related set. These included nsp2, nsp7 and nsp13 (**Fig. 4a**). Our algorithm identified cadherin 2, α-catenin, β-catenin, δ-catenin, and ZO 1 and 2 as the most susceptible proteins to SARS-CoV-2 infection (**Fig. 4b**).

**Fig. 4.**
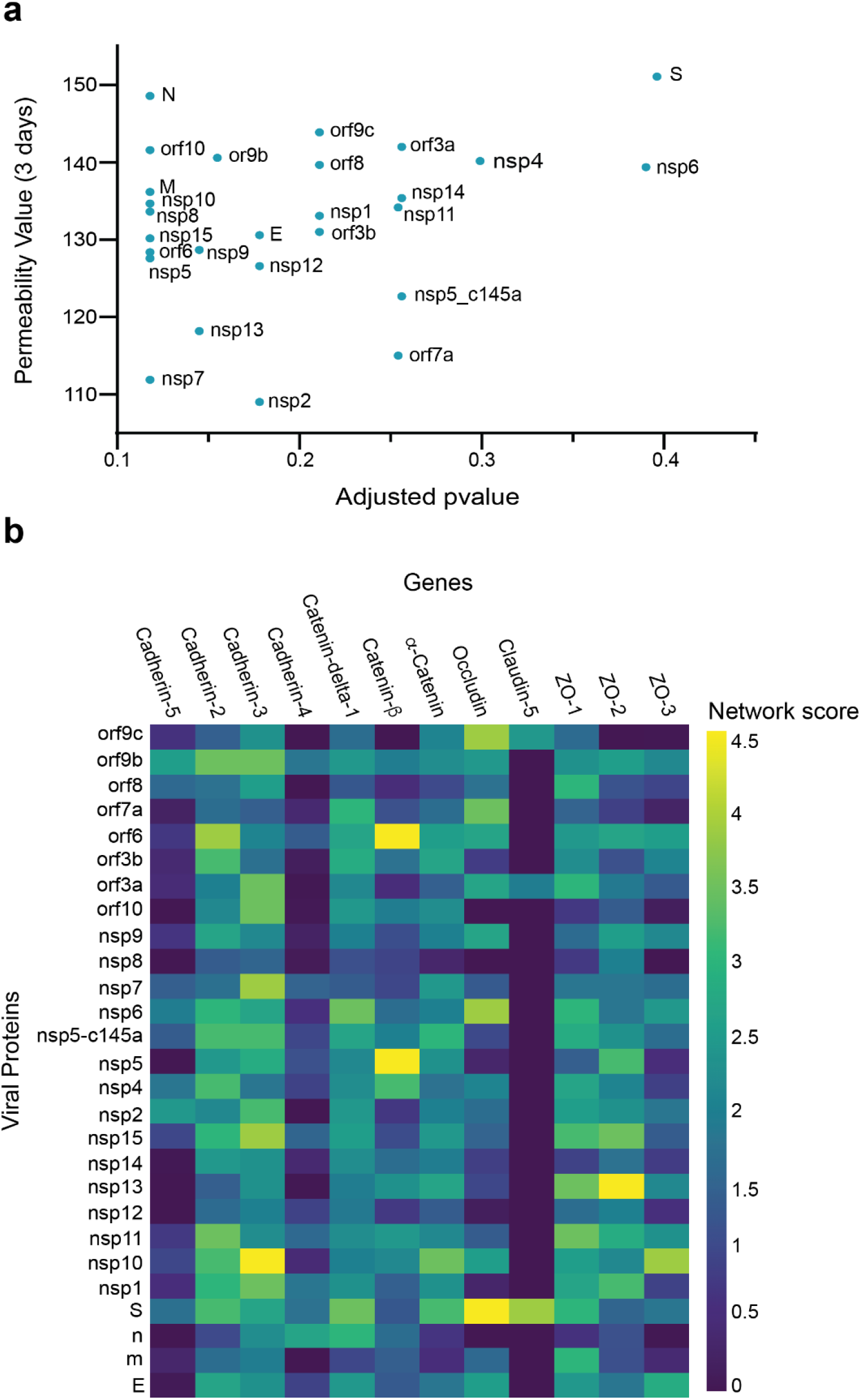
Correlation between viral protein effect on permeability and proximity to permeability-related proteins in a PPI network. **a** Correlation of adjusted *p*-value versus permeability (Pearson = 0.295); **b** proximity between vascular proteins and the viral proteins.

As our network propagation model was highly correlated with our experimental results, we applied it to other physiological systems that are known to be affected by SARS-CoV-2. We created a list of all proteins that are known to be affected by the SARS-CoV-2 proteins according to the literature (**SI Table 1 white columns)**. The table was composed of both proteins that were identified experimentally via western blot, proteomics, and IHC (marked in blue) and those identified clinically as being highly correlated with loss of specific functionality in specific tissues (marked in red). We then applied the network-based model to identify which of the proteins in **SI Table 1** are most susceptible to the different SARS-CoV-2 proteins. As can be seen in **Fig. 5a**, **SI Tables 1 and 2,** and **SI Figs. 1–7**, specific SARS-CoV-2 proteins were identified as affecting specific proteins in specific tissues. As expected, most of the SARS-CoV-2 proteins affected more than one protein, the most salient being nsp11, nsp4 and nsp7 (**Fig. 5b**), each of which was predicted to affect more than 40 different proteins. An additional parameter that should be taken into account is the protein’s “distance” from the viral proteins. This value represents the number of hops in the PPI network from a given protein to the viral proteins, where a value of 1 represents a direct viral–host connection. We hypothesized that the closer the distance between the viral proteins and the given protein, the more significant the viral effect. **SI Table 1 (gray columns)** and **Fig. 5c** present the calculated distances. Most of the proteins that were identified in **SI Table 1** were classified with a distance of 1 or 2 from the virus, suggesting more severe putative effects.

**Fig. 5.**
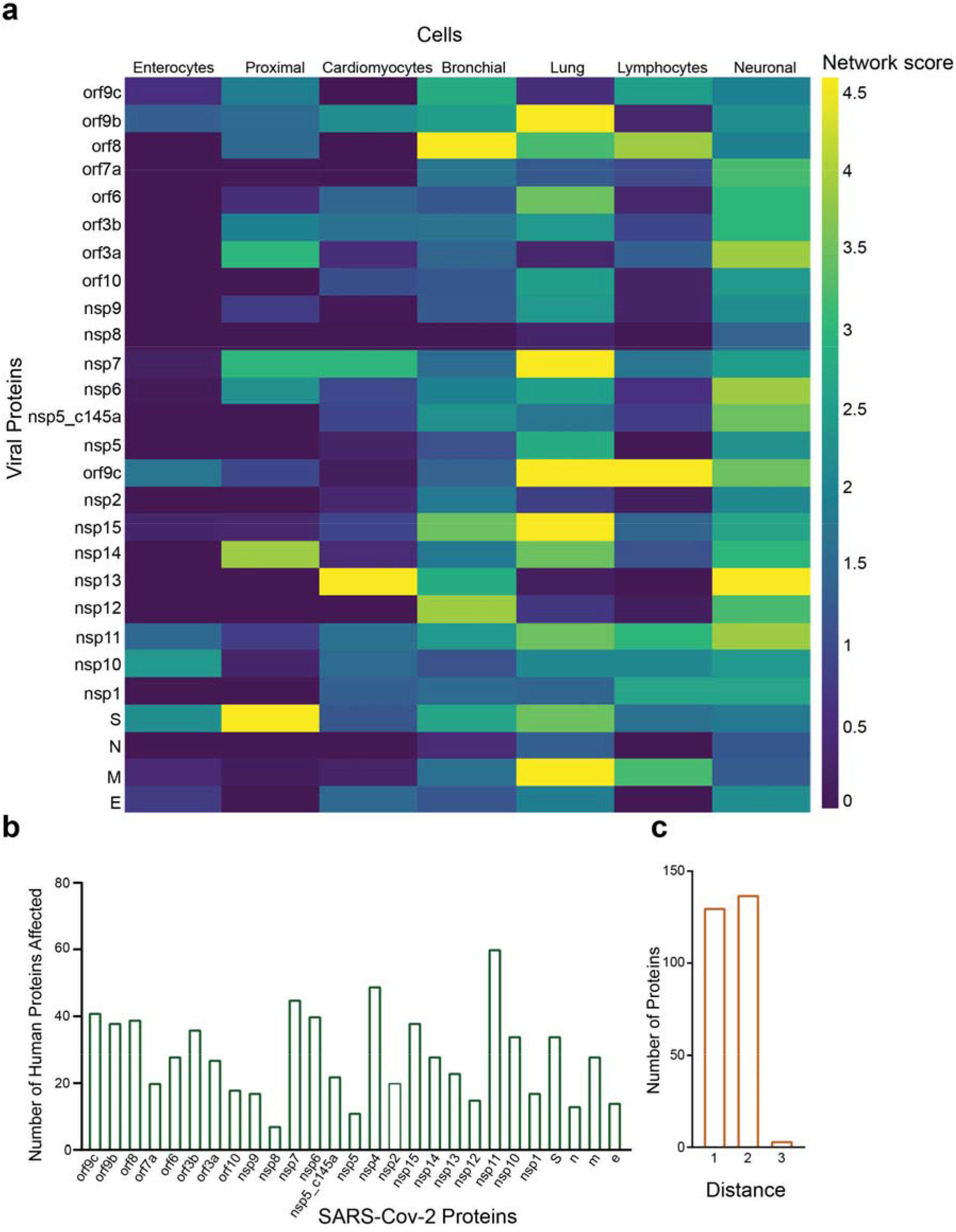
Protein identification using PPI. **a** PPI results for the SARS-CoV-2 proteins that have a significant effect on the proteins presented in SI Table 1 for each system; **b** number of proteins affected by each SARS-CoV-2 protein, as calculated by PPI; **c** number of proteins with a specific distance factor from the viral proteins (also shown in SI Table 1).

## Discussion

Due to the impact of SARS-CoV-2, many studies have looked at the physiological responses the virus^1–4,19^. In this work, we sought to identify how specific SARS-CoV-2 proteins affect the vasculature by assessing the effect of individual SARS-CoV-2 proteins on endothelial cells (HUVEC). There are major advantages to this approach: it enables pinpointing and isolating how each of the SARS-CoV-2 proteins independently affects the endothelial response, and measuring endothelial functionality directly.

The current study showed that almost 70% (18 out of 26) of the SARS-CoV-2 proteins affect endothelial permeability; however, the most significant proteins were nsp2, nsp5_c145a and nsp7, which also induced upregulated expression of the coagulation factor VWF and cytokine release. These important facts can shed light on the multiple pathologies observed in SARS-CoV-2 infection, which include cytokine storm, increased coagulation, increased cardiovascular disease, increased neurological symptoms, and a significant increase in coagulation-related diseases (e.g., heart attack and stroke)^1,5^. The results presented here demonstrate how the vasculature becomes leaky, which can cause exotoxicity, i.e., the penetration of toxic reagents from the blood into the brain. While there are many parameters associated with functional changes, the use of advanced tools, including network-based analysis, enabled us to elucidate the specific proteins and the specific interactions that are predicted to cause these changes. The PPI network enabled us to predict that the changes observed in barrier function are probably due to interactions between host proteins such cadherin 2, α-catenin, β-catenin, δ-catenin, and ZO 1 and 2, and the viral proteins nsp2, nsp5_c145a and nsp7.

PPI analysis revealed a highly correlated effect of nsp7 and nsp13 on β-catenin in endothelial cells (**Fig. 4b**). As the inhibition of wnt/β-catenin signaling found in multiple sclerosis impairs the blood– brain barrier and increases the infiltration of immune cells, it is reasonable to assume that a similar mechanism is caused by SARS-CoV-19, i.e., damage to endothelial cells that leads to increased permeability and leakage of blood vessels, infiltration of immune cells, and activation of an immune reaction that results in damage to the infected organ^33,34^. Interestingly, neither nsp2 nor nsp5_c145a affected a high number of proteins (**Fig. 5b**), whereas nsp7 did, as identified by the network. Analyzing the repertoire of SARS-CoV-2 proteins, we see almost no effect of the structural proteins; rather, mostly nonstructural and open reading frame proteins affected HUVEC functionality, manifested as decreased barrier function and increased cytokine secretion (**Figs. 2** and **3**). While the nonstructural proteins are mainly responsible for the replication of viral RNA, the open reading frame proteins are related to counteraction with the host immune system; some of these are localized to the mitochondria and have been shown to alter the mitochondrial antiviral signaling pathway^35^. We found that the proteins most affecting barrier function (decreased TEER and CD31 expression) and cytokine response (IL-6 secretion and VWF expression) were nsp2, nsp5_c145a and nsp7 (**Figs. 2** and **3**); nsp7 forms a replication complex with nsp8 and nsp12 that is essential for viral replication and transcription^24,36^. Peng et al.^37^ suggested that in the core polymerase complex nsp7–nsp8–nsp12, nsp12 is the catalytic subunit, and nsp7 and nsp8 function as cofactors. They further suggested that the mechanism of activation mainly involves the cofactors rather than the catalytic subunit^37^. This might explain why we saw mainly an effect of the cofactor proteins on endothelial cells and almost no effect of the catalytic subunit. Network interactions^38^ have shown that nsp7 has the most interactions with the host, suggesting a potential target for treatment of COVID-19. Moreover, no mutations were found in nsp7 compared to nsp2 or nsp5_c145a^39^, suggesting a conserved protein with a vital function in virus survival. The nsp13 protein has both helicase activity and 5’ triphosphatase activity, which play an important role in mRNA capping. We saw a large effect of nsp13 on barrier function, but hardly any effect on cytokine secretion. Chen et al.^40^ suggested functional complexation between nsp8 and nsp12, the RdRp (RNA dependent-RNA polymerase) replication complex, and nsp13. Given the fact that we observed a very large effect of nsp7—one of the proteins of the replication complex—and an effect of nsp13 on HUVEC barrier function, complexation of nsp13 with the replication complex might indicate an important role for this complex in the impaired functionality of the HUVEC, and therefore in the propagation of the disease, and the known vascular damage seen in COVID-19 patients. This may also position nsp13 as an important protein affecting all cell types (**Fig. 5a**), and a target for disease treatment. Many studies have looked at the SARS-CoV-2 interaction with nonpulmonary/nonvascular tissues (e.g., neurons, hepatocytes, immune components such as lymphocytes, macrophages, etc.)^1^, as pathological studies identified a viral effect on these tissues, despite their very limited amount, or lack of ACE2 receptors. To better understand how SARS-CoV-2 interacts with and affects other tissues, we consolidated all of the proteins that are currently known to be affected by the virus into **SI Table 1**. It is interesting to note that the most dominant SARS-CoV-2 proteins are nsp4, nsp11 and nsp7. Davies et al.^41^ identified the interaction of nsp2 with nsp4, both involved in endoplasmic reticulum (ER) calcium signaling and mitochondrial biogenesis. This suggests a new functional role in the host ER and mitochondrial organelle contact process and calcium homeostasis.

By now it is clear that vasculature plays a significant role in the physiological response to the virus. However, it is still unclear how the virus affects the vasculature, and if it can be found in the blood. This is a very important question, as it has significant consequences on the extent of the virus’s ability to affect the vasculature. Current studies demonstrate that the pulmonary vasculature is significantly affected and is one of the dominant triggers for the aforementioned pathologies. However, involvement with the rest of the vasculature is still unclear, as is whether the virus can be found in an active form in the blood circulation^36,42–45^. Some studies suggest that even if there are traces of SARS-CoV-2 in the blood, it is not in an active form, and cannot cause disease or a systemic response^44^. On the other hand, some studies suggest that SARS-CoV-2 can be found in the blood, and can induce the disease and cause both cellular and systemic dysfunction^36,42,46^. While this question is beyond the scope of this work, it is important to note that if future studies do identify the active form of SARS-CoV-2 in human blood, then the implications of our findings will apply to this systemic response as well^47,48^.

As already noted, the pathology is probably a combination of multiple conditions and pathways which are activated by the different proteins. However, our findings might open new avenues for future therapeutics. Moreover, most of the proteins that were identified as affected by SARS-CoV-2 had a distance factor of at most 3 to the human and viral proteins. This coincides with the current dogma, whereby proteins that have a shorter distance between them are more likely to be affected.

Finally, we would like to point out some of the limitations of our study. The two major limitations of our approach are: **(a)** inability to identify the effect of multiple proteins; **(b)** neglecting the effect of the coronavirus structure and binding on the cellular response. The former point can be overcome by combining different SARS-CoV-2 proteins in a well. However, since the SARS-CoV-2 expresses 29 proteins, there are 29! combinations, which is about ~9 × 10^30^. Therefore, we decided to focus on individual proteins, and allow further studies to pursue any combinations of interest. Regarding the latter limitation, we did not include the coronavirus structure (including the ACE2 receptors) in this study, because many studies have already demonstrated the cellular response to this structure^19,49,50^, and how tissues that do not have significant ACE2 expression (such as neurons, immune components such as B and T lymphocytes, and macrophages) are affected by the virus remains an open question.

## Conclusions

Accumulating clinical evidence suggests that COVID-19 is a vascular disease. However, only a few studies have identified the specific role of each of the SARS-CoV-2 proteins in the cellular response. In this work, we characterized the endothelial response to each of 26 SARS-CoV-2 proteins and identified those that have the most significant effect on the barrier function. In addition, we used PPI network-based analysis to predict which of the endothelial proteins is most affected by the virus and to identify the specific role of each of the SARS-CoV-2 proteins in the observed changes in systemic protein expression. Overall, this work identified which of the SARS-CoV-2 proteins are most dominant in their effect on the physiological response to the virus. We believe that the data presented in this work will give us better insight into the mechanism by which the vasculature and the system respond to the virus, and will enable us to expedite drug development for the virus by targeting the identified dominant proteins.

## Methods

### Generation of Lentiviral SARS-CoV-2 Constructs

Plasmids encoding the SARS-CoV-2 open reading frames proteins and eGFP control were a kind gift of Nevan Krogan (Addgene plasmid #141367-141395). Plasmids were acquired as bacterial LB–agar stabs, and used per the provider’s instructions. Briefly, each stab was first seeded in LB agar (Bacto Agar; BD (Biosciences, San Jose, CA)) in 10-cm plates. Then, single colonies were inoculated into flasks containing LB (BD Difco LB Broth, Lennox) and 100 μg/ml penicillin (Biological Industries, Beit HaEmek, Israel). Transfection-grade plasmid DNA was isolated from each flask using the ZymoPURE II Plasmid Maxiprep Kit (Zymo Research, Irvine, CA) according to the manufacturer’s instructions. HEK293T cells (ATCC, Manassas, VA) were seeded in 10-cm cell-culture plates at a density of 4 × 10^6^ cells/plate. The cells were maintained in 293T medium composed of DMEM high glucose (4.5 g/l; Merck, NJ) supplemented with 10% fetal bovine serum (FBS; Biological Industries), 1X NEAA (Biological Industries), and 2 mM L-alanine–L-glutamine (Biological Industries, Israel). The following day, the cells were transfected with a SARS-CoV-2 orf-expressing plasmid and the packaging plasmids using TransIT-LT1 transfection reagent (Mirus Bio, Madison, WI) according to the provider’s instructions. Briefly, 6.65 μg SARS-CoV-2 lentivector plasmid, 3.3 μg pVSV-G (vesicular stomatitis virus G protein), and 5 μg psPAX2 were mixed in Opti-MEM reduced serum medium (Gibco, (Waltham, MA)) with 45 μl of TransIT-LT1, kept at room temperature (RT) to form a complex, and then added to each plate. Following 18 h of incubation, the transfection medium was replaced with 293T medium and virus-rich supernatant was harvested after 48 h and 96 h. The supernatant was clarified by centrifugation (500*g*, 5 min) and filtration (0.45 μm, Millex-HV, Merck Millipore (Burlington, MA)). All virus stocks were aliquoted and stored at −80 °C.

### Lentivirus preparation

Lentiviral stocks, pseudo-typed with VSV-G, were produced in HEK293T cells as previously described^51^. Briefly, each of the pLVX plasmids containing the SARS-CoV-2 genes were cotransfected with third-generation lentivirus helper plasmids at equimolar ratio; 48 h later, the lentivirus-containing medium was collected for subsequent use.

### Endothelial Cell Culture

HUVEC (PromoCell GmbH, Heidelberg, Germany) were used to test each viral protein’s impact on vascular properties. After thawing, the HUVEC were expanded in low-serum endothelial cell growth medium (PromoCell) at 37 °C with 5% CO_2_ in a humidifying incubator, and used at passage p4–p6. Cells were grown to 80–90% confluence before being transferred to transparent polyethylene terephthalate (PET) Transwell supports (0.4 um pore size, Greiner Bio-One, Austria) or a glass-bottom well plate (Cellvis(Mountain View, CA)). Before seeding, the uncoated substrates were treated with Entactin-Collagen IV-Laminin (ECL) Cell Attachment Matrix (Merck) diluted in DMEM (10 μg/cm^2^) for 1 h in the incubator. Then, the HUVEC, harvested using a DetachKit (PromoCell), were seeded inside the culture platforms at a density of 250,000 cells/cm^2^ and grown for 3 days. Then viral infection with the different plasmids was performed and its impact on cell behavior was tested 3 days later.

### TEER measurement

The barrier properties of the endothelial monolayer were evaluated by TEER measurements, 3 and 4 days after viral infection. TEER was measured with the Millicell ERS-2 Voltohmmeter (Merck Millipore). TEER values (Ω cm^2^) were calculated and compared to those obtained in a Transwell insert without cells, considered as a blank, in three different individual experiments, with two inserts used for each viral protein.

### Immunofluorescence

HUVEC plated on glass-bottom plates were rinsed in phosphate buffered saline (PBS) and fixed in 4% paraformaldehyde (Sigma-Aldrich, Rehovot, Israel) for 20 min at RT, 5 days after viral infection. Immunocytochemistry was carried out after permeabilization with 0.1% Triton X-100 (Sigma-Aldrich, Rehovot, Israel) in PBS for 10 min at RT and blocking for 30 min with 5% FBS in PBS. The following primary antibodies were applied overnight in PBS at 4 °C: rabbit anti-VWF (Abcam, Cambridge, UK) and rabbit anti-CD31 (Abcam) against platelet endothelial cell adhesion molecule 1 (PECAM1). Cells were then washed three times in PBS and stained with the secondary antibody, anti-rabbit Alexa Fluor 488 (Invitrogen, Carlsbad, CA), for 1 h at RT. After four washes with PBS, cells were incubated with Hoechst in PBS for 10 min at RT to stain the nuclei. After two washes with PBS, imaging was carried out using an inverted confocal microscope (Olympus FV3000-IX83) with suitable filter cubes and equipped with 20× (0.8 NA) and 60× (1.42 NA) objectives. Image reconstruction and analysis were done using open-source ImageJ software^52^.

### Network analysis

We scored the effect of each viral protein on selected human proteins using network propagation^25^. Specifically, a viral protein was represented by the set of its human interactors^23^; each of these received a prior score, equal to 1/n, where n is the size of the interactor set; these scores were propagated in a network of protein–protein interactions^53^. To assess the statistical significance of the obtained scores, we compared them to those computed on 100 randomized networks that preserve node degrees.

### Quantitative ELISA for IL-6

ELISA was performed on conditioned medium of infected HUVEC 3 days postinfection, according to the manufacturer’s recommendations (PeproTech Rehovot, Israel).

### Statistical analysis

The results are presented as mean ± SD, unless otherwise indicated. Statistically significant differences among multiple groups were evaluated by F-statistic with two-way ANOVA, followed by the Holm–Sidak test for multiple comparisons (GraphPad Prism 8.4.3). The difference between the two data sets was assessed and *p* < 0.05 was considered statistically significant.

### Data availability

All of the data supporting the findings of this study are available from the corresponding authors upon reasonable request.

## Acknowledgements

B.M.M. was supported by the Azrieli Foundation, Israel Science Foundation (ISF grant: 2248/19), ERC SweetBrain 851765, TEVA and Aufzien Center for Prevention of Parkinson (APPD). U.A. was supported by the Israel Science Foundation (ISF grant 953/16), the German Research Foundation (DFG) (NA: 207/10-1) and the Taube/Koret Global Collaboration in Neurodegenerative Diseases. R.S. was supported by the Israel Science Foundation (ISF grant 2417/20), within the Israel Precision Medicine Partership program. The work of Y.N., A.E. and K.I. was supported by European Research Council Consolidator Grant OCLD (project no. 681870).

## Competing interests

There are no conflicts to declare

## Author contributions

### Additional information

**Supplementary information** includes:

Supplementary Figures 1–7.

Supplementary Tables 1 and 2

## Notes

### Competing Interest Statement

The authors have declared no competing interest.

