## Supplementary Information for "Effect of SARS-CoV-2 proteins on vascular permeability"

### Supplementary Information
### Effect of SARS-CoV-2 proteins on vascular permeability

Rossana Rauti[1*], Meishar Shahoha[2,3*], Yael Leichtmann-Bardoogo[1 *], Rami Nasser[4], Rina Tamir[1], Victoria Miller[1], Tal Babich[1,2], Kfir Shaked [1,2], Avner Ehrlich[5], Konstantinos Ioannidis[5], Yaakov Nahmias [5], Roded Sharan[4], Uri Ashery [2,3,6#], and Ben M. Maoz[1,3,6#]

[1]Department of Biomedical Engineering, Tel Aviv University, Israel
[2]School of Neurobiology, Biochemistry and Biophysics, The George S. Wise Faculty of Life Sciences, Tel Aviv University, Tel Aviv, Israel
[3]Sagol School of Neuroscience, Tel Aviv University, Tel Aviv, Israel
[4]Blavatnik School of Computer Science, Tel Aviv University, Tel Aviv, Israel
[5]Hebrew University of Jerusalem - Grass Center for Bioengineering
[6]The Center for Nanoscience and Nanotechnology, Tel Aviv University, Israel

[*] These authors are equal contributors
### Shared corresponding authors

Corresponding author
Uri Ashery, mail:
Ben M. Maoz, mail:

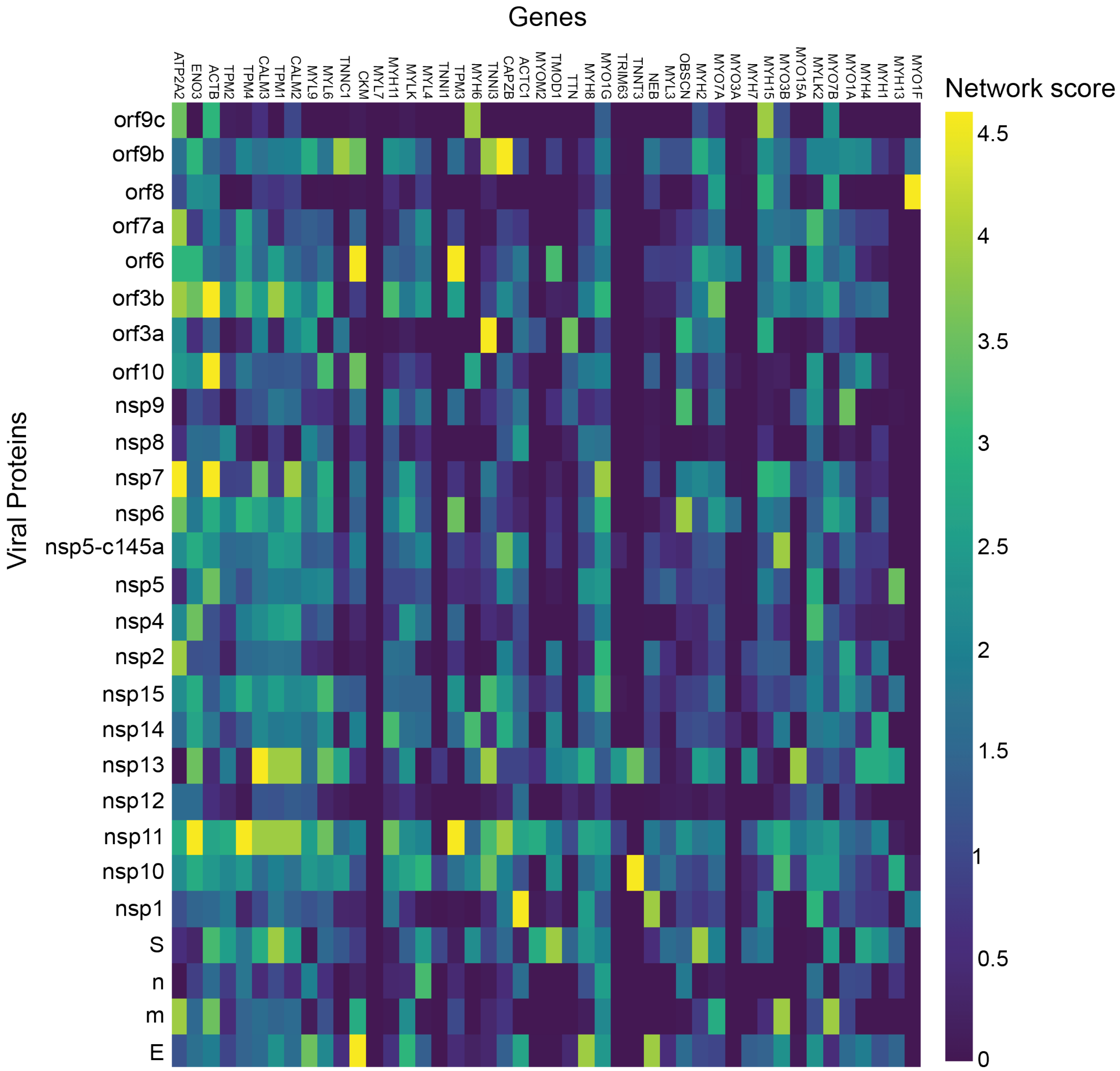


**SI Fig 1. Cardiomyocytes.** pvalue correlation of a target protein with a specific viral protein, calculated empirically using 100 random samples.

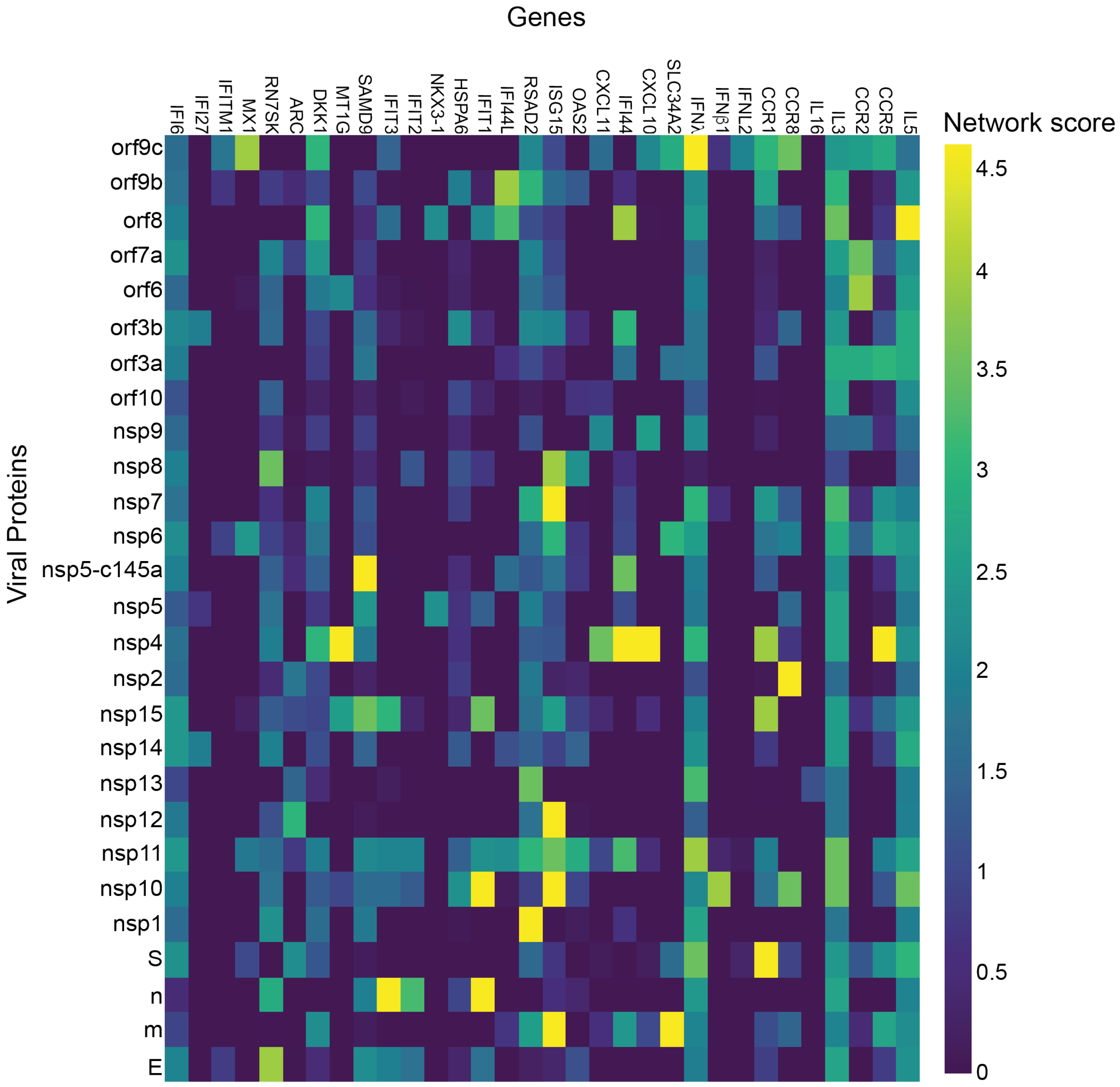


**SI Fig 2. Enterocytes.** pvalue correlation of a target protein with a specific viral protein, calculated empirically using 100 random samples.

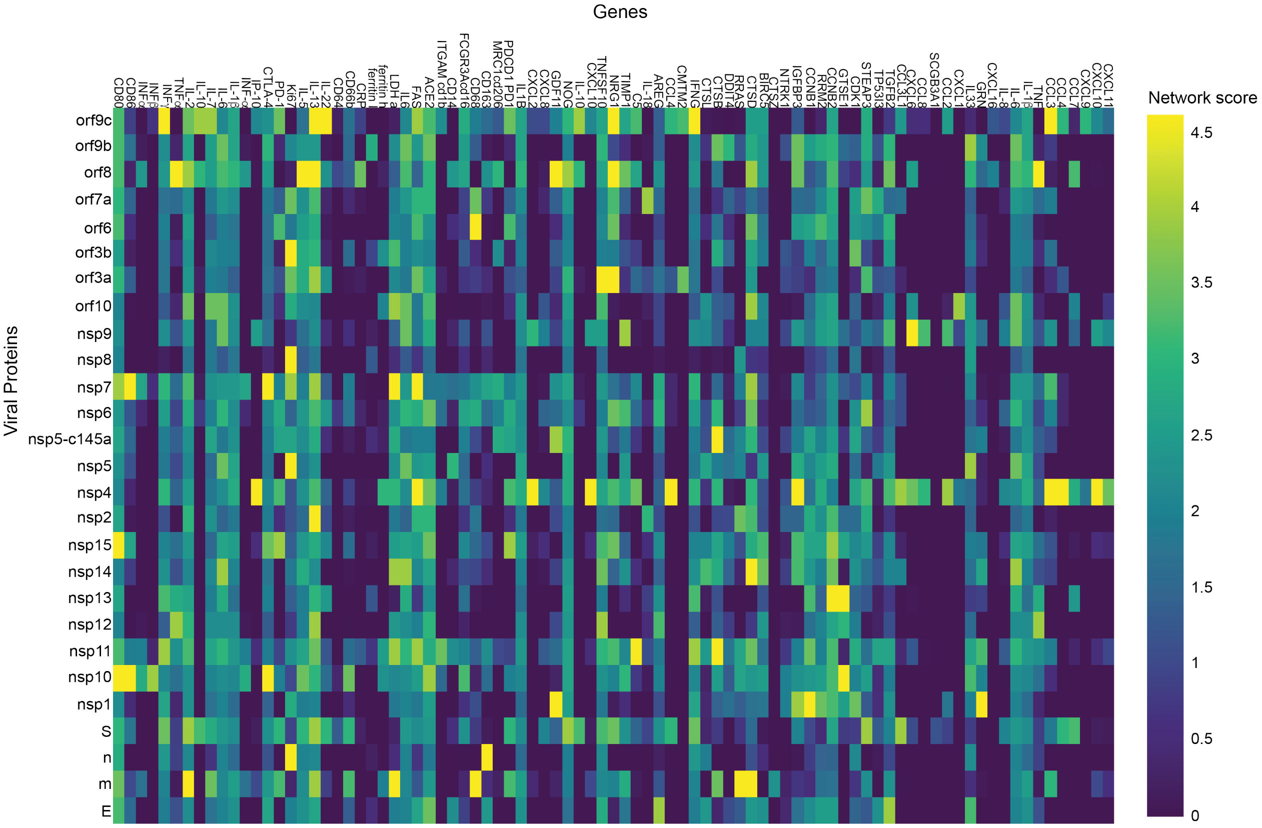


**SI Fig 3. Lymphocytes.** pvalue correlation of a target protein with a specific viral protein, calculated empirically using 100 random samples.

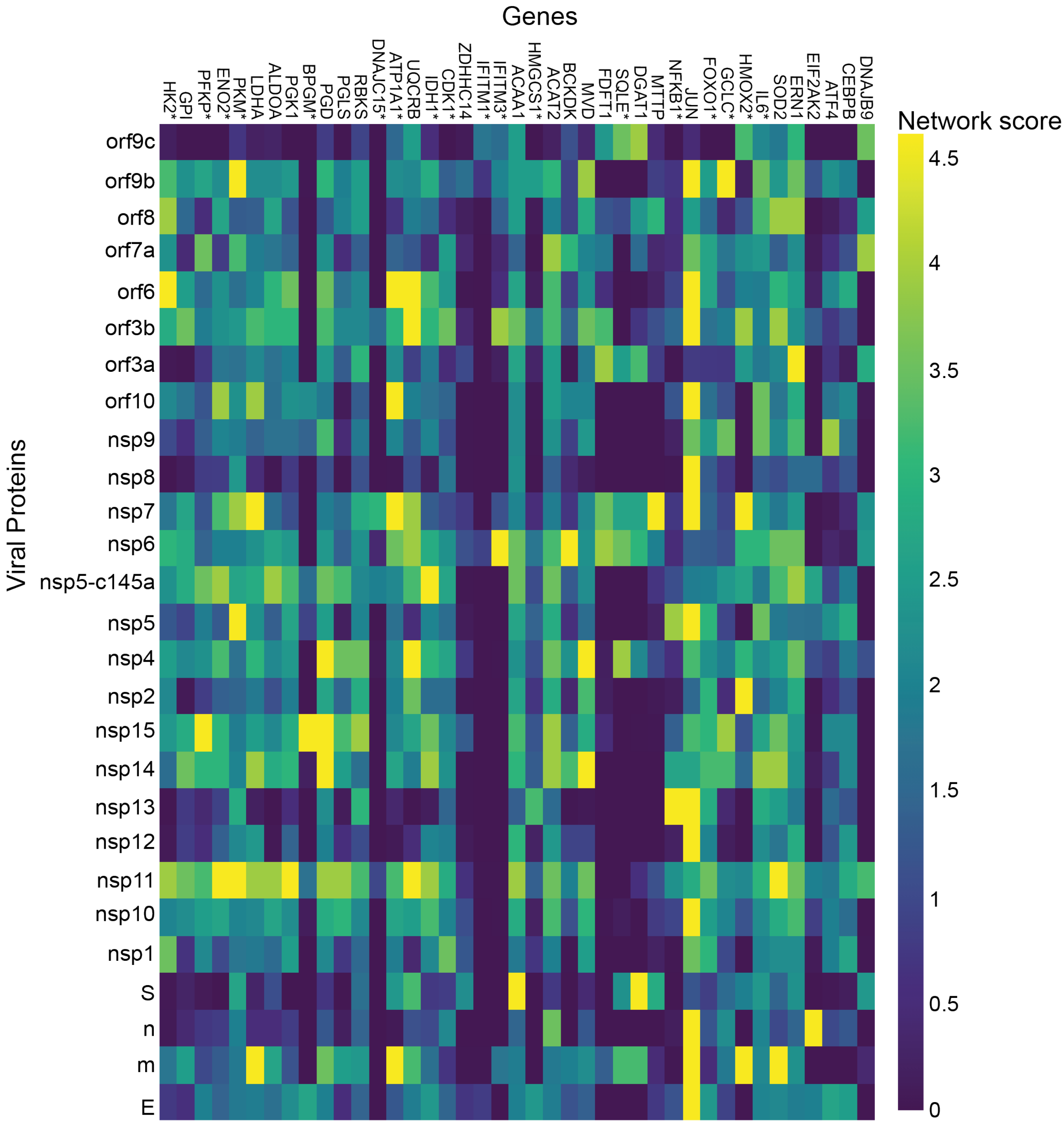


**SI Fig 4. Lung.** pvalue correlation of a target protein with a specific viral protein, calculated empirically using 100 random samples.

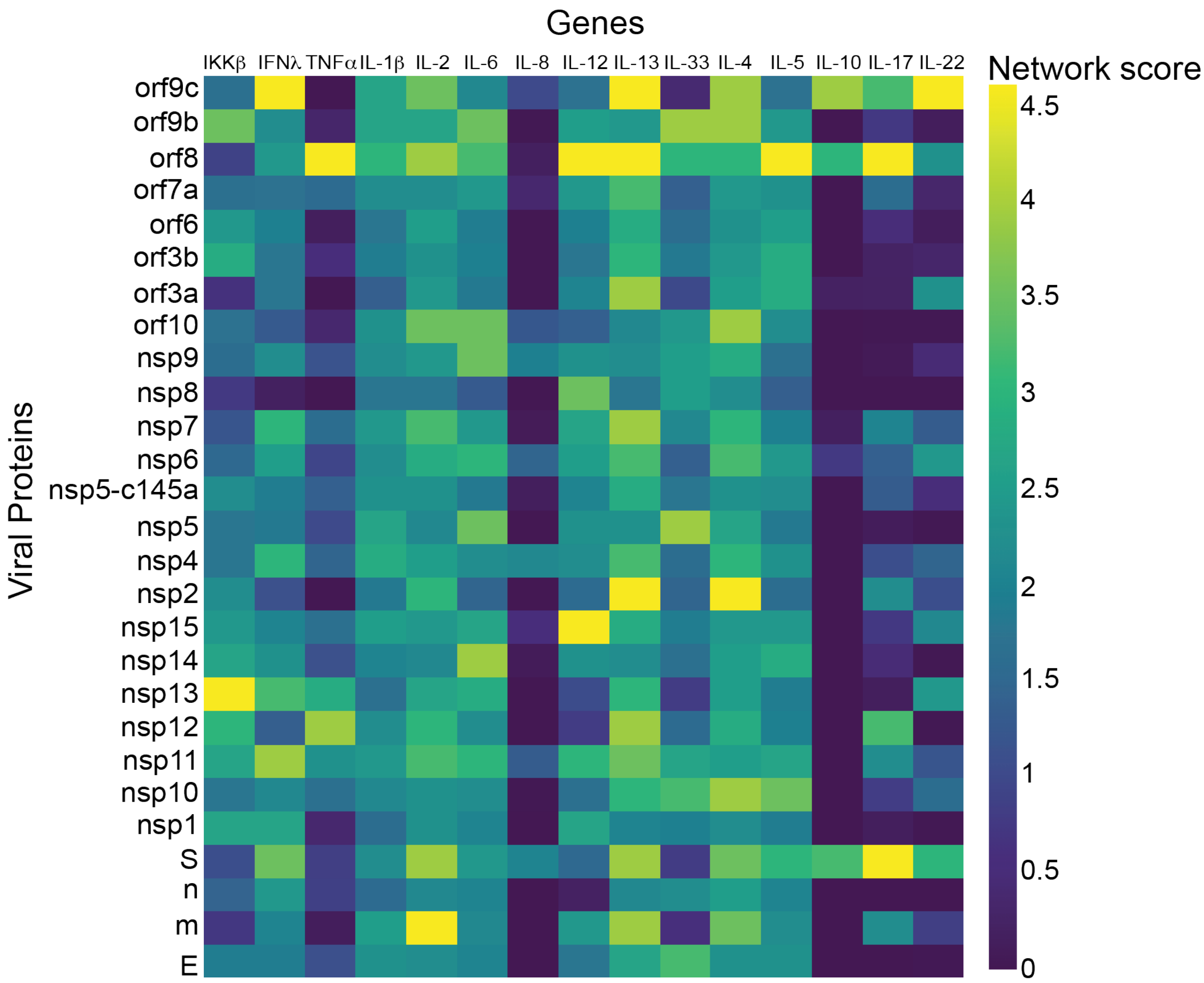


**SI Fig 5. Bronchial.** pvalue correlation of a target protein with a specific viral protein, calculated empirically using 100 random samples.

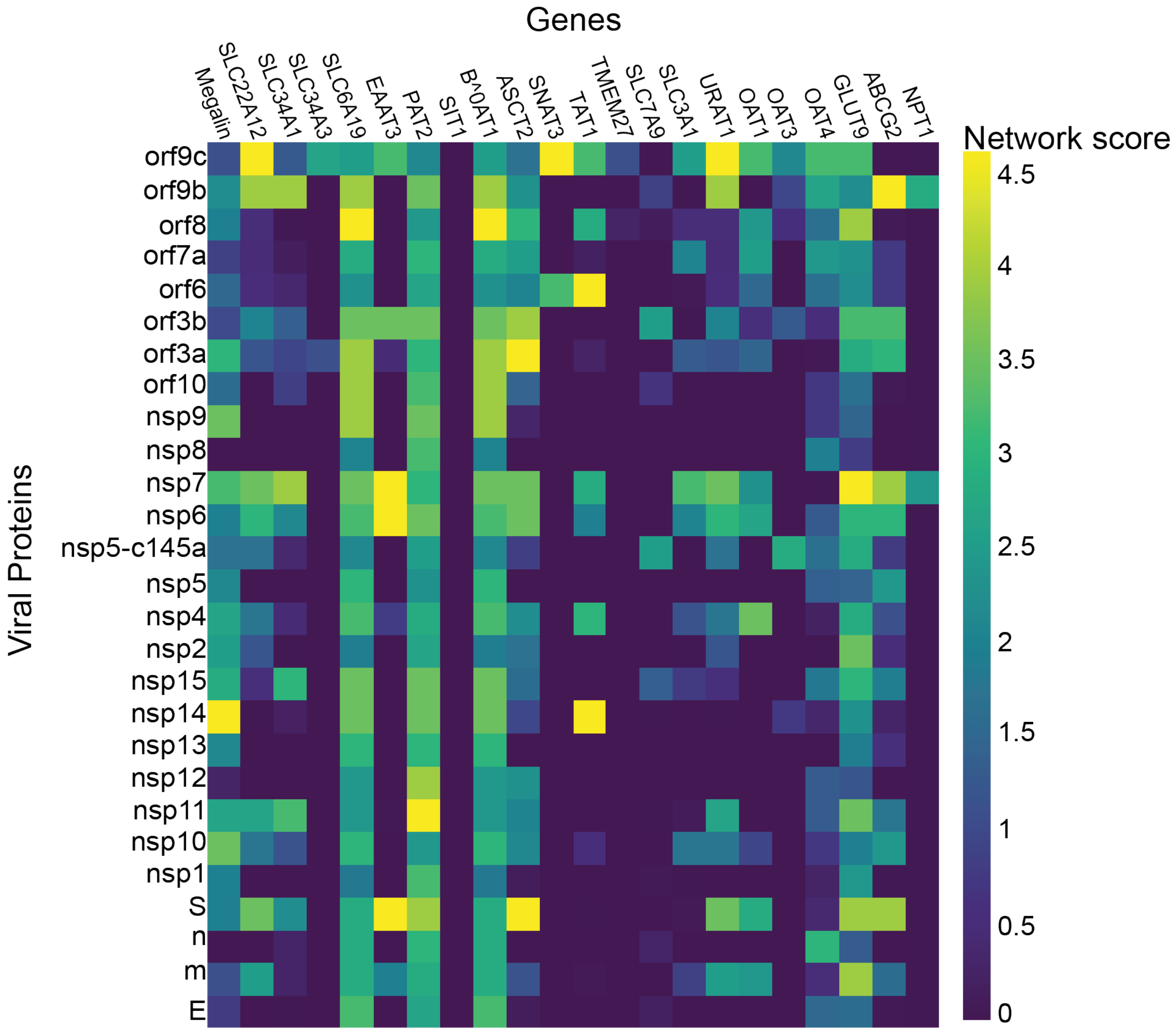


**SI Fig 6. Proximal tubule cells.** value correlation of a target protein with a specific viral protein, calculated empirically using 100 random samples.

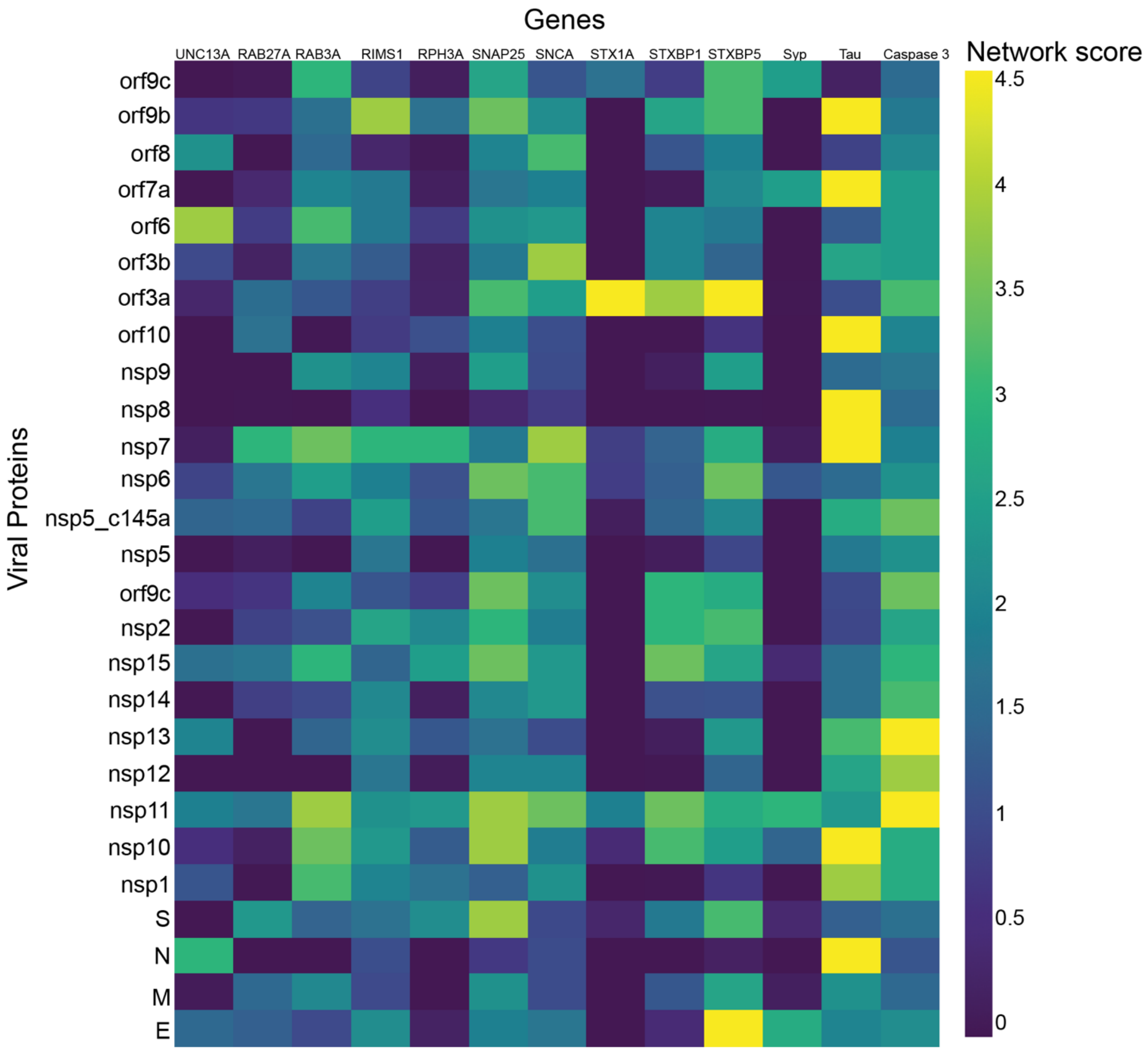


**SI Fig 7. Neuronal cells.** value correlation of a target protein with a specific viral protein, calculated empirically using 100 random samples.

**SI Table 1. Documented change by SARS-Cov-2.**

| Cell type | Function | Gene | Entrez Gene | Identified significant SARS-Cov-19 proteins | Distance |
|---|---|---|---|---|---|
| Dendritic cells (T cell proliferation) | Impaired dendritic cells, induces T-cell proliferation and cytokine production and signal for activation of the T-cell | CD80[1] | 941 | [e; m; spike; nsp10; nsp11; nsp15; nsp7; orf3a; orf6; orf7a; orf8; orf9b; orf9c] | 2 |
| | | CD86[1] | 942 | [nsp10; nsp7] | 2 |
| | | INFα[1] | 3439 | | 2 |
| | | INF β[1] | 3456 | | 2 |
| T cell | Impaired T cell maturation and activation, reduced proliferation | INF λ[1-4] | 3458 | | 2 |
| | | TNFα[1,3,4] | 7124 | [nsp12; orf8] | 1 |
| | Production of pro-inflammatory cytokines | IL-2[2,3,4] | 3558 | [m; spike; nsp11; nsp12; nsp2; nsp7; orf10; orf8; orf9c] | 2 |
| | | IL-10[2,3,4] | 3586 | [spike; orf8; orf9c] | 2 |
| | | IL-7[2] | 3574 | [m; orf10; orf9c] | 2 |
| | | IL-6[2,4] | 3569 | [nsp11; nsp14; nsp5; nsp6; nsp9; orf10; orf8; orf9b] | 2 |

| | | | | | |
|---|---|---|---|---|---|
| | | IL-1β[2] | 3553 | [orf8] | 2 |
| | | INF-α[2] | 3439 | | 2 |
| | | MOP-1[2] | | | |
| | | IP-10[2] | 3627 | | 2 |
| | reduced proliferation | CTLA-4[3] | 1493 | [nsp10; nsp15; nsp7; orf9b] | 1 |
| | | PD-1[1,3] | 5133 | [m; spike; nsp15; nsp4; orf6; orf9c] | 1 |
| | | Ki67[3] | 4288 | [n; nsp5; nsp7; nsp8; orf3a; orf3b; orf7a] | 1 |
| | | IL-5[3] | 3567 | [spike; nsp10; orf8] | 2 |
| | | IL-13[3] | 3596 | [m; spike; nsp10; nsp11; nsp12; nsp13; nsp2; nsp4; nsp6; nsp7; orf3a; orf3b; orf7a; orf8; orf9c] | 2 |
| | | IL-9[3,4] | 3578 | | |
| | | IL-22[3] | 50616 | [spike; orf9c] | 3 |
| NK | Decreased **IFN-γ production** by NK cells | INFλ[5] | 3458 | | 2 |
| Myeloid cells | systemic inflammation may impair the myeloid cells functionality | HLA-DR[6] | | | |
| | | CD64[6] | 2209 | | 2 |

| | | | | | |
|---|---|---|---|---|---|
| | | CD66b[6] | 1088 | [m; spike; nsp10] | 2 |
| | | CRP[6] | 1401 | [orf8] | 1 |
| | | Ferritin l[6] | 2512 | | 1 |
| | | ferritin h[6] | 2495 | [nsp4] | 1 |
| | | D-dimer[6] | | | |
| | | fibrinogen[7] | | | |
| | | LDH a[6] | 3939 | [m; nsp11; nsp14; nsp4; nsp7; orf10; orf3b] | 1 |
| | | IL6[6] | 3569 | [nsp11; nsp14; nsp5; nsp6; nsp9; orf10; orf8; orf9b] | 2 |
| Lymphocyte | severe lymphocyte apoptosis | FAS[7] | 355 | [nsp11; nsp2; nsp4; nsp7; orf3a; orf6; orf7a; orf9c] | 1 |
| Monocyte and macrophage | Activation and transcription of proinflammatory genes | ACE2[4] | 59272 | [e; nsp10; nsp12; nsp15; nsp2; nsp4; nsp6; orf10; orf7a; orf8; orf9b; orf9c] | 1 |
| | | IL6[4] | 3569 | [nsp11; nsp14; nsp5; nsp6; nsp9; orf10; orf8; orf9b] | 2 |
| | | IL-10[4] | 3586 | [spike; orf8; orf9c] | 2 |
| | | TNFα[4] | 7124 | [nsp12; orf8] | 1 |
| | | ITGAM cd1b[4] | 3684 | [nsp11] | 2 |

| | | | | | |
|---|---|---|---|---|---|
| | | CD14[4] | 929 | [nsp5] | 2 |
| | | FCGR3Acd16[4] | 2214 | [nsp10] | 2 |
| | | CD68[4] | 968 | [m; orf6; orf7a] | 1 |
| | | CD80[4] | 941 | [e; m; spike; nsp10; nsp11; nsp15; nsp7; orf3a; orf6; orf7a; orf8; orf9b; orf9c] | 2 |
| | | CD163[4] | 9332 | [n] | 1 |
| | | MRC1 cd206[4] | 4360 | [orf8] | 2 |
| | | PDCD1 PD1[4] | 5133 | | 1 |
| PB mononuclear cells | Activation of proinflammatory genes | IL1B[4] | 3553 | [orf8] | 2 |
| | | CXCL2[4] | 2920 | [nsp4; nsp9] | 2 |
| | | CXCL8[4] | 3576 | | 2 |
| PBMC | Activation of proinflammatory genes | GDF11[8] | 10220 | [nsp1; nsp5_c145a; orf8] | 1 |
| | | NOG[8] | 9241 | [spike; nsp5_c145a; orf3a; orf8; orf9c] | 2 |
| | | IL10[8] | 3586 | [spike; orf8; orf9c] | 2 |
| | | CXCL10[8] | 3627 | nsp4 | 2 |

| | | | | | |
|---|---|---|---|---|---|
| | | TNFSF10[8] | 8743 | [spike; nsp11; nsp12; nsp14; nsp15; orf10; orf3a; orf9b] | 1 |
| | | NRG1[8] | 3084 | [spike; nsp11; nsp15; nsp4; nsp6; orf3a; orf6; orf7a; orf8; orf9c] | 1 |
| | | TIMP1[8] | 7076 | [nsp10; nsp6; nsp9; orf8] | 2 |
| | | C5[8] | 727 | [nsp11; nsp4; orf9c] | 2 |
| | | IL18[8] | 3606 | [nsp2; orf7a] | 2 |
| | | AREG[8] | 374 | [e; nsp12; nsp6] | 2 |
| | | CCL4[8] | 6351 | [spike; nsp4; orf9c] | 1 |
| | | CMTM2[8] | 146225 | [orf3a; orf9c] | 1 |
| | | CXCL8[8] | 3576 | | 2 |
| | | IFNλ[8] | 3458 | [spike; nsp11; nsp13; nsp4; nsp7; orf9c] | 2 |
| Lymphocytes | SARS-CoV-2 infection may cause lymphocyte apoptosis | CTSL[8] | 1514 | [nsp14] | 1 |
| | | CTSB[8] | 1508 | [m; nsp11; nsp4; nsp5_c145a; nsp6; orf9b] | 1 |
| | | DDIT4[8] | 54541 | [orf9b] | 2 |

| | | | | | |
|---|---|---|---|---|---|
| | | RRAS[8] | 6237 | [m; nsp2] | 1 |
| | | CTSD[8] | 1509 | [m; nsp11; nsp14; nsp2; nsp4; nsp5_c145a; nsp6; nsp7; orf10; orf6; orf8] | 1 |
| | | BIRC5[8] | 332 | [nsp14; nsp15] | 1 |
| | | TNFSF10[8] | 8743 | [spike; nsp11; nsp12; nsp14; nsp15; orf10; orf3a; orf9b] | 1 |
| | | CTSZ[8] | 1522 | | 1 |
| | | NTRK1[8] | 4914 | | 1 |
| | | IGFBP3[8] | 3486 | [nsp1; nsp14; nsp15; nsp4; orf8; orf9b] | 1 |
| | | CCNB1[8] | 891 | [nsp1; nsp11; nsp13] | 1 |
| | | RRM2[8] | 6241 | [nsp1; orf6] | 2 |
| | | CCNB2[8] | 9133 | [nsp1; nsp10; nsp11; nsp13; nsp14; nsp15; nsp2; orf8; orf9b] | 1 |
| | | GTSE1[8] | 51512 | [nsp10; nsp12; nsp13] | 1 |
| | | CDK1[8] | 983 | [nsp1; orf3b] | 1 |
| | | STEAP3[8] | 55240 | [spike; nsp4; nsp6; orf3a; orf7a] | 2 |

| | | | | | |
|---|---|---|---|---|---|
| | | TP53I3[8] | 9540 | | 2 |
| BALF | Activation of proinflammatory cytokines | CXCL2[8] | 2920 | [nsp4; nsp9] | 2 |
| | | TGFB2[8] | 7042 | [e; nsp4; nsp9; orf6; orf8; orf9b] | 2 |
| | | CCL3L1[8] | 6349 | [spike; nsp4] | 2 |
| | | CXCL6[8] | 6372 | [nsp4; nsp9] | 2 |
| | | CCL8[8] | 6355 | [nsp4; nsp9] | 2 |
| | | SCGB3A1[8] | 92304 | | 2 |
| | | TNFSF10[8] | 8743 | [spike; nsp11; nsp12; nsp14; nsp15; orf10; orf3a; orf9b] | 1 |
| | | CCL2[8,9] | 6347 | [nsp4; nsp9] | 2 |
| | | CXCL1[8] | 2919 | [orf10] | 1 |
| | | IL33[8] | 90865 | [e; nsp10; nsp5; orf8; orf9b] | 2 |
| | | GRN[8] | 2896 | [nsp1; nsp11] | 1 |
| | | CXCL16[8] | 58191 | | 1 |
| Immune cells of BALF | higher levels of inflammatory cytokines in lung macrophages from patients with severe COVID-19 infection. | IL-8[9] | 3576 | | 2 |

| | | | | | |
|---|---|---|---|---|---|
| | | IL-6[9] | 3569 | [nsp11; nsp14; nsp5; nsp6; nsp9; orf10; orf8; orf9b] | 2 |
| | | IL-1β[9] | 3553 | [orf8] | 2 |
| | | TNFα[9] | 7124 | [nsp12; orf8] | 1 |
| | | CCL3[9] | 6348 | [m; nsp4; nsp7; orf9c] | 1 |
| | | CCL4[9] | 6351 | [spike; nsp4; orf9c] | 1 |
| | | CCL7[9] | 6354 | [spike; orf8] | 1 |
| | | CXCL9[9] | 4283 | [orf9c] | 1 |
| | | CXCL10[9] | 3627 | [nsp4] | 2 |
| | | CXCL11[9] | 6373 | [nsp4] | 2 |
| Primary lung epithelial cells | glycolysis | HK2*[10] | 3099 | [nsp1; nsp11; nsp6; orf6; orf8; orf9b] | 1 |
| | glycolysis | GPI[10] | 2821 | [nsp11; nsp14; orf3] | 1 |
| | glycolysis | PFKP[10] | 5214 | [nsp11; nsp14; nsp15; nsp5_c145a; orf7a] | 1 |
| | glycolysis | ENO2*[10] | 2026 | [nsp11; nsp14; nsp15; nsp4; nsp5_c145a; nsp7; orf10] | 1 |

| | | | | | |
|---|---|---|---|---|---|
| | glycolysis | PKM*[10] | 5315 | [nsp11; nsp5; nsp7; orf7a; orf9b] | 1 |
| | glycolysis | LDHA[10] | 3939 | [m; nsp11; nsp14; nsp4; nsp7; orf10; orf3b] | 1 |
| | glycolysis | ALDOA[10] | 226 | [nsp11; nsp5_c145a; orf3b; orf6] | 1 |
| | glycolysis | PGK1[10] | 5230 | [nsp11; nsp14; nsp5_c145a; orf3b; orf6] | 1 |
| | glycolysis | BPGM*[10] | 669 | [nsp15] | 2 |
| | Pentose-Phosphate | PGD[10] | 5226 | [m; nsp11; nsp14; nsp15; nsp4; nsp5_c145a; nsp9; orf3b; orf6; orf9b] | 1 |
| | Pentose-Phosphate | PGLS[10] | 25796 | [nsp10; nsp11; nsp15; nsp4; nsp5_c145a] | 1 |
| | Pentose-Phosphate | RBKS[10] | 64080 | [nsp11; nsp13; nsp15; nsp4; orf3a] | 2 |
| | mitochondria | DNAJC15*[10] | 29103 | [nsp7] | 2 |
| | mitochondria | ATP1A1*[10] | 476 | [m; nsp11; nsp6; nsp7; orf10; orf3b; orf6] | 1 |
| | mitochondria | UQCRB[10] | 7381 | [m; spike; nsp11; nsp2; nsp4; nsp5_c145a; nsp6; nsp7; orf3b; orf6] | 1 |

| | | | | | |
|---|---|---|---|---|---|
| | mitochondria | IDH1*[10] | 3417 | [nsp10; nsp11; nsp14; nsp15; nsp4; nsp5_c145a; orf3b; orf6] | 1 |
| | mitochondria | CDK1*[10] | 983 | [nsp1; orf3b] | 1 |
| | palmitoylation | ZDHHC14[10] | 79683 | | 2 |
| | palmitoylation | IFITM1*[10] | 8519 | | 2 |
| | palmitoylation | IFITM3*[10] | 10410 | [nsp6; orf3b] | 1 |
| | lipogenesis | ACAA1[10] | 30 | [spike; nsp11; nsp12; nsp14; nsp15; nsp5_c145a; nsp6; orf3b] | 1 |
| | lipogenesis | HMGCS1*[10] | 3158 | [nsp13] | 1 |
| | lipogenesis | ACAT2[10] | 39 | [n; nsp10; nsp11; nsp14; nsp15; nsp4; nsp5_c145a; nsp6; orf3b; orf6; orf7a; orf9b] | 1 |
| | lipogenesis | BCKDK[10] | 10295 | [nsp14; nsp6; orf7a] | 0 |
| | lipogenesis | MVD[10] | 4597 | [nsp10; nsp11; nsp14; nsp2; nsp4; orf3b; orf9b] | 2 |
| | lipogenesis | FDFT1[10] | 2222 | [nsp6; nsp7; orf3a; orf3b] | 1 |
| | lipogenesis | SQLE*[10] | 6713 | [m; nsp4; nsp6; orf9c] | 2 |
| | lipogenesis | DGAT1[10] | 8694 | [m; spike; nsp6; orf3a; orf9c] | 2 |

| | | | | | |
|---|---|---|---|---|---|
| | lipogenesis | MTTP[10] | 4547 | [nsp7; orf8] | 2 |
| | oxidative stress | NFKB1*[10] | 4790 | [nsp13; nsp5] | 1 |
| | oxidative stress | JUN[10] | 3725 | [e; m; n; nsp1; nsp10; nsp12; nsp13; nsp15; nsp4; nsp5; nsp7; nsp8; nsp9; orf10; orf3b; orf6; orf7a; orf9b] | 1 |
| | oxidative stress | FOXO1*[10] | 2308 | [nsp1; nsp11; nsp14; nsp15; nsp2; nsp5] | 1 |
| | oxidative stress | GCLC*[10] | 2729 | [nsp14; nsp15; nsp9; orf9b] | 1 |
| | oxidative stress | HMOX2*[10] | 3163 | [m; nsp2; nsp4; nsp6; nsp7; orf3b; orf9c] | 1 |
| | oxidative stress | IL6*[10] | 3569 | [nsp11; nsp14; nsp5; nsp6; nsp9; orf10; orf8; orf9b] | 2 |
| | oxidative stress | SOD2[10] | 6648 | [m; nsp11; nsp14; nsp15; nsp5_c145a; nsp6; orf3b; orf6; orf8] | 1 |
| | ER stress | ERN1[10] | 2081 | [nsp10; nsp11; nsp4; nsp5_c145a; nsp6; nsp9; orf3a; orf8; orf9b; orf9c] | 1 |
| | ER stress | EIF2AK2[10] | 5610 | [n] | 1 |
| | ER stress | ATF4[10] | 468 | [nsp9] | 1 |
| | ER stress | CEBPB[10] | 1051 | | 2 |

| | | | | | |
|---|---|---|---|---|---|
| | ER stress | DNAJB9[10] | 4189 | [nsp11; orf7a; orf9c] | 2 |
| Cardiomyocytes | cardiac muscle tissue development<br>regulation of viral process<br>actin filament organization<br>proteosomal protein catabolic process<br>cellular respiration<br>regulation of lymphocyte activation<br>sensory perception of smell<br>pos. reg. of receptor sig.pathway via JAK-STAT | ATP2A2[11] | 488 | [m; nsp2; nsp6; nsp7; orf3b; orf6; orf7a; orf9c] | 1 |
| | | ENO3[11] | 2027 | [nsp11; nsp13; nsp4; orf3b; orf6; orf9b] | 1 |
| | | ACTB[11] | 60 | [m; spike; nsp5; nsp7; orf10; orf3b; orf9c] | 1 |
| | | TPM2[11] | 7169 | | 1 |
| | | TPM4[11] | 7171 | [nsp11; nsp6; orf3b] | 1 |
| | | CALM3[11] | 808 | [nsp11; nsp13; nsp7] | 1 |
| | | TPM1[11] | 7168 | [spike; nsp11; nsp13; orf3b] | 1 |
| | | CALM2[11] | 805 | [nsp11; nsp13; nsp7] | 1 |
| | | MYL9[11] | 10398 | [e] | 1 |

| | | | | | |
|---|---|---|---|---|---|
| | | MYL6[11] | 4637 | [nsp11; nsp13; nsp15; orf10; orf3b] | 1 |
| | | TNNC1[11] | 7134 | [orf9b] | 2 |
| | | CKM[11] | 1158 | [e; orf10; orf6; orf9b] | 1 |
| | | MYL7[11] | 58498 | | 2 |
| | | MYH11[11] | 4629 | [nsp11; nsp14; orf3b] | 1 |
| | | MYLK[11] | 4638 | [e] | 2 |
| | | MYL4[11] | 4635 | [n; nsp10] | 2 |
| | | TNNI1[11] | 7135 | | 2 |
| | | TPM3[11] | 7170 | [nsp11; nsp6; orf6] | 1 |
| | | MYH6[11] | 4624 | [nsp14; orf9c] | 1 |
| | | TNNI3[11] | 7137 | [nsp10; nsp11; nsp13; nsp15; orf3a; orf9b] | 1 |
| | | CAPZB[11] | 832 | [nsp11; nsp5_c145a; orf9b] | 1 |
| | | ACTC1[11] | 70 | [nsp1] | 1 |
| | | MYOM2[11] | 9172 | | 2 |
| | | TMOD1[11] | 7111 | [spike; orf6] | 1 |

| | | | | | |
|---|---|---|---|---|---|
| | | TTN[11] | 7273 | [orf3a] | 1 |
| | | MYH8[11] | 4626 | [e] | 1 |
| | | MYO1G[11] | 64005 | [nsp15; nsp2; nsp6; nsp7; orf3b] | 1 |
| | | TRIM63[11] | 84676 | | 1 |
| | | TNNT3[11] | 7140 | [nsp10; nsp13] | 1 |
| | | NEB[11] | 4703 | [e; nsp1] | 2 |
| | | MYL3[11] | 4634 | | 2 |
| | | OBSCN[11] | 84033 | [nsp6; nsp9; orf3a] | 1 |
| | | MYH2[11] | 4620 | [spike] | 1 |
| | | MYO7A[11] | 4647 | [orf3b] | 2 |
| | | MYO3A[11] | 53904 | | 2 |
| | | MYH7[11] | 4625 | | 1 |
| | | MYH15[11] | 22989 | [nsp7; orf8; orf9c] | 2 |
| | | MYO3B[11] | 140469 | [m; nsp10; nsp5_c145a] | 1 |
| | | MYO1H[11] | 283446 | | 1 |
| | | MYO15A[11] | 51168 | [nsp13] | 2 |
| | | MYLK2[11] | 85366 | [nsp1; nsp4; orf7a] | 2 |
| | | MYO7B[11] | 4648 | [m] | 2 |

| | | | | | |
|---|---|---|---|---|---|
| | | MYO1A[11] | 4640 | [nsp9] | 1 |
| | | MYH4[11] | 4622 | | 1 |
| | | MYH1[11] | 4619 | | 1 |
| | | MYH13[11] | 8735 | [nsp5] | 1 |
| | | MYO1F[11] | 4542 | [orf8] | 1 |
| Human bronchial epithelial cell line (16HBE) | Activation of proinflammatory cytokines | IKKβ, [12] | 3551 | [nsp12; nsp13; orf9b] | 1 |
| | | IFNλ[12] | 3458 | [spike; nsp11; nsp13; nsp4; nsp7; orf9c] | 2 |
| | | TNF-α[12] | 7124 | [nsp12; orf8] | 1 |
| | | LTa3[12] | | | |
| | | GMCSF[12] | | | |
| | | IL-1β[12] | 3553 | [orf8] | 2 |
| | | IL-2[12] | 3558 | [m; spike; nsp11; nsp12; nsp2; nsp7; orf10; orf8; orf9c] | 2 |
| | | IL-6[12] | 3569 | [nsp11; nsp14; nsp5; nsp6; nsp9; orf10; orf8; orf9b] | 2 |
| | | IL-8[12] | 3576 | | 2 |
| | | IL-12[12] | 3593 | [nsp11; nsp15; nsp8; orf8] | 1 |

| | | | | | |
|---|---|---|---|---|---|
| | | IL-13[12] | 3596 | [m; spike; nsp10; nsp11; nsp12; nsp13; nsp2; nsp4; nsp6; nsp7; orf3a; orf3b; orf7a; orf8; orf9c] | 2 |
| | | IL-33[12] | 90865 | [e; nsp10; nsp5; orf8; orf9b] | 2 |
| | | IL-4[12] | 3565 | [m; spike; nsp10; nsp2; nsp4; nsp6; nsp7; orf10; orf8; orf9b; orf9c] | 2 |
| | | IL-5[12] | 3567 | [spike; nsp10; orf8] | 2 |
| | | IL-10[12] | 3586 | [spike; orf8; orf9c] | 2 |
| | | IL-17[12] | 3605 | [spike; nsp12; orf8; orf9c] | 1 |
| | | IL-22[12] | 50616 | [spike; orf9c] | 3 |
| Neuronal cells | altered distribution of Tau | Tau[13] | 4137 | [n; nsp1; nsp10; nsp13; nsp7; nsp8; orf10; orf7a; orf9b] | 1 |
| | apoptosis of neuronal cells | Caspase 3[13] | 836 | [nsp11; nsp12; nsp13; nsp14; nsp15; nsp4; nsp5_c145a; orf3a] | 1 |
| | | UNC13A | 23025 | [n; orf6] | 1 |
| | | RAB27A | 5873 | [nsp7] | 2 |
| | | RAB3A | 5864 | [nsp1; nsp10; nsp11; nsp15; nsp7; orf6; orf9c] | 1 |
| | | RIMS1 | 22999 | [nsp7; orf9b] | 2 |

| | | | | | |
|---|---|---|---|---|---|
| | | RPH3A | 22895 | [nsp7] | 1 |
| | | SNAP25 | 6616 | [spike; nsp10; nsp11; nsp15; nsp2; nsp4; nsp6; orf3a; orf9b] | 2 |
| | | SNCA | 6622 | [nsp11; nsp5_c145a; nsp6; nsp7; orf3b; orf8] | 1 |
| | | STX1A | 6804 | [orf3a] | 1 |
| | | STXBP1 | 6812 | [nsp10; nsp11; nsp15; nsp2; nsp4; orf3a] | 1 |
| | | STXBP5 | 134957 | [e; spike; nsp2; nsp6; orf3a; orf9b; orf9c] | 2 |
| | | Syp | 6855 | [nsp11] | 2 |
| Proximal tubule cells | decreased expression of the multi-ligand receptor megalin LRP2 | Megalin[14] | 4036 | [nsp10; nsp14; nsp7; nsp9; orf3a] | 1 |
| | decreased expression of the multi-ligand receptor megalin LRP3 | SLC22A12[14] | 116085 | [spike; nsp6; nsp7; orf9b; orf9c] | 2 |
| | decreased expression of the multi-ligand receptor megalin LRP4 | SLC34A1[14] | 6569 | [nsp11; nsp15; nsp7; orf9b] | 2 |

| | | | | | |
|---|---|---|---|---|---|
| | decreased expression of the multi-ligand receptor megalin LRP5 | SLC34A3[14] | 142680 | | 2 |
| | decreased expression of the multi-ligand receptor megalin LRP6 | SLC6A19[14] | 340024 | [e; nsp10; nsp13; nsp14; nsp15; nsp4; nsp5; nsp6; nsp7; nsp9; orf10; orf3a; orf3b; orf8; orf9b] | 2 |
| | amino acid transporter | EAAT3[14] | 6505 | [spike; nsp6; nsp7; orf3b; orf9c] | 1 |
| | amino acid transporter | PAT2[14] | 153201 | [n; spike; nsp1; nsp11; nsp12; nsp13; nsp14; nsp15; nsp6; nsp7; nsp8; nsp9; orf10; orf3a; orf3b; orf7a; orf9b] | 2 |
| | amino acid transporter | SIT1[14] | 54716 | | 2 |
| | amino acid transporter | B^0AT1[14] | 340024 | | 2 |
| | amino acid transporter | B^0AT3[14] | 348932 | | 1 |
| | amino acid transporter | ASCT2[14] | 6510 | [spike; nsp6; nsp7; orf3a; orf3b; orf8] | 2 |
| | amino acid transporter | SNAT3[14] | 10991 | [orf6; orf9c] | 2 |
| | amino acid transporter | TAT1[14] | 117247 | [nsp14; nsp4; orf6; orf9c] | 2 |
| | amino acid transporter | TMEM27[14] | 57393 | | 2 |
| | amino acid transporter | SLC7A9[14] | 11136 | | 2 |
| | amino acid transporter | SLC3A1[14] | 6519 | [nsp7] | 2 |

| | | | | | |
|---|---|---|---|---|---|
| | impaired tubular handling of uric acid | URAT1[14] | 116085 | | 2 |
| | impaired tubular handling of uric acid | OAT1[14] | 9356 | [nsp4; orf9c] | 2 |
| | impaired tubular handling of uric acid | OAT3[14] | 9376 | | 2 |
| | impaired tubular handling of uric acid | OAT4[14] | 55867 | [n; orf9c] | 2 |
| | impaired tubular handling of uric acid | GLUT9[14] | 56606 | [m; spike; nsp11; nsp15; nsp2; nsp6; nsp7; orf3b; orf8; orf9c] | 1 |
| | impaired tubular handling of uric acid | ABCG2[14] | 9429 | [spike; nsp6; nsp7; orf3a; orf3b; orf9b] | 2 |
| | impaired tubular handling of uric acid | NPT1[14] | 6568 | | 1 |
| | impaired tubular handling of uric acid | NPT4[14] | 10786 | | 2 |
| Enterocytes (gut organoids) | Defense response to virus | IFI6[15] | 2537 | | 2 |
| | | IFI27[15] | 3429 | | 2 |
| | | IFITM1[15] | 8519 | | 2 |
| | | MX1[15] | 4599 | [orf9c] | 1 |
| | | RN7SK[15] | 125050 | [e; nsp8] | 1 |
| | | ARC[15] | 23237 | [nsp12] | 2 |
| | | DKK1[15] | 22943 | [nsp4; orf8; orf9c] | 2 |
| | | MT1G[15] | 4495 | [nsp4] | 2 |

| | | | | | |
|---|---|---|---|---|---|
| | | SAMD9[15] | 54809 | [nsp15; nsp5_c145a] | 2 |
| | | IFIT3[15] | 3437 | [n; nsp15] | 1 |
| | | C10orf99[15] | 387695 | | 1 |
| | | IFIT2[15] | 3433 | [n] | 1 |
| | | NKX3-1[15] | 4824 | | 1 |
| | | LINC00941[15] | 1E+08 | | |
| | | HSPA6[15] | 3310 | | 1 |
| | | IFIT1[15] | 3434 | [n; nsp10; nsp15] | 1 |
| | | IFI44L[15] | 10964 | [orf8; orf9b] | 2 |
| | | CMPK2[15] | 129607 | | 1 |
| | | RSAD2[15] | 91543 | [nsp1; nsp11; nsp13; orf9b] | 2 |
| | | ISG15[15] | 9636 | [m; nsp10; nsp11; nsp12; nsp6; nsp7; nsp8] | 2 |
| | | OAS2[15] | 4939 | | 2 |
| | | CXCL11[15] | 6373 | [nsp4] | 2 |
| | | IFI44[15] | 10561 | [nsp11; nsp4; nsp5_c145a; orf3b; orf8] | 1 |
| | | CXCL10[15,16] | 3627 | [nsp4] | 2 |
| | | SLC34A2[15] | 10568 | [m; nsp6] | 2 |

| | | | | | |
|---|---|---|---|---|---|
| | | IFNλ[16,17] | 3458 | [spike; nsp11; nsp13; nsp4; nsp7; orf9c] | 3 |
| | | IFN b1[17] | 3456 | [nsp10] | 2 |
| | | IFNL2[16] | 282616 | | 2 |
| | | IFNL3[16] | 28617 | | 1 |
| | | CCR1[16] | 1230 | [spike; nsp15; nsp4; orf9c] | 2 |
| | | CCR8[16] | 1237 | [nsp10; nsp2; orf9c] | 2 |
| | | IL16[16] | 3603 | | 2 |
| | | IL3[16] | 3562 | [nsp10; nsp11; nsp7; orf8; orf9b] | 2 |
| | | CCR2[16] | 729230 | [orf6; orf7a] | 2 |
| | | CCR5[16] | 1234 | [nsp4; orf3a] | 2 |
| | | IL5[16] | 3567 | [spike; nsp10; orf8] | 2 |

**SI Table 2. Significant target for each viral protein.**

| SARS-Cov-2 Proteins | Human proteins which are most effect | Number of proteins |
|---|---|---|
| **e** | [RN7SK; SLC6A19; MYL9; CKM; MYLK; MYH8; NEB; IL-33; JUN; CD80; ACE2; AREG; TGFB2; STXBP5] | 14 |
| m | [ISG15; SLC34A2; GLUT9; ATP2A2; ACTB; MYO3B; MYO7B; IL-2; IL-13; IL-4; LDHA; PGD; ATP1A1*; UQCRB; SQLE*; DGAT1; JUN; HMOX2*; SOD2; CD80; IL-7; PD-1; CD66b; CD68; CTSB; RRAS; CTSD; CCL3] | 28 |
| n | [IFIT3; IFIT2; IFIT1; PAT2; OAT4; MYL4; ACAT2; JUN; EIF2AK2; Ki67; CD163; UNC13A; Tau] | 13 |
| S | [IFNλ; CCR1; IL-5; SLC22A12; EAAT3; PAT2; ASCT2; GLUT9; ABCG2; ACTB; TPM1; TMOD1; MYH2; IL-2; IL-13; IL-4; IL-10; IL-17; IL-22; UQCRB; ACAA1; DGAT1; CD80; PD-1; CD66b; NOG; TNFSF10; NRG1; CCL4; STEAP3; CCL3L1; CCL7; SNAP25; STXBP5] | 34 |
| nsp1 | [RSAD2; PAT2; ACTC1; NEB; MYLK2; HK2*; CDK1*; JUN; FOXO1*; GDF11; IGFBP3; CCNB1; RRM2; CCNB2; GRN; RAB3A; Tau] | 17 |
| nsp10 | [IFIT1; ISG15; IFN b1; CCR8; IL-3; IL-5; Megalin; SLC6A19; MYL4; TNNI3; TNNT3; MYO3B; IL-13; IL-33; IL-4; PGLS; IDH1*; ACAT2; MVD; JUN; ERN1; CD80; CD86; CTLA-4; CD66b; ACE2; FCGR3Acd16; TIMP1; CCNB2; GTSE1; RAB3A; SNAP25; STXBP1; Tau] | 34 |
| nsp11 | [RSAD2; ISG15; IFI44; IFNλ; IL-3; SLC34A1; PAT2; GLUT9; ENO3; TPM4; CALM3; TPM1; CALM2; MYL6; MYH11; TPM3; TNNI3; CAPZB; IL-2; IL-6; IL-12; IL-13; HK2*; GPI; PFKP; ENO2*; PKM*; LDHA; ALDOA; PGK1; PGD; PGLS; RBKS; ATP1A1*; UQCRB; IDH1*; ACAA1; ACAT2; MVD; FOXO1*; SOD2; ERN1; DNAJB9; CD80; FAS; ITGAM cd1b; TNFSF10; NRG1; C5; CTSB; CTSD; CCNB1; CCNB2; GRN; RAB3A; SNAP25; SNCA; STXBP1; Syp; Caspase 3] | 61 |
| nsp12 | [ARC; ISG15; PAT2; IKKβ; TNF-α; IL-2; IL-13; IL-17; ACAA1; JUN; ACE2; TNFSF1; AREG; GTSE1; Caspase 3] | 15 |
| nsp13 | [RSAD2; IFNλ; SLC6A19; PAT2; ENO3; CALM3; TPM1; CALM2; MYL6; TNNI3; TNNT3; MYO15A; IKKβ; IL-13; RBKS; HMGCS1*; NFKB1*; JUN; CCNB1; CCNB2; GTSE1; Tau; Caspase 3] | 23 |

| | | |
|---|---|---|
| nsp14 | [Megalin; SLC6A19; PAT2; TAT1; MYH11; MYH6; IL-6; GPI; PFKP; ENO2*; LDHA; PGK1; PGD; IDH1*; ACAA1; ACAT2; BCKDK; MVD; FOXO1*; GCLC*; SOD2; TNFSF10'; CTSL; CTSD; BIRC5; IGFBP3; CCNB2; Caspase 3] | 28 |
| nsp15 | [SAMD9; IFIT3; IFIT1; CCR1; SLC34A1; SLC6A19; PAT2; GLUT9; MYL6; TNNI3; MYO1G; IL-12; PFKP; ENO2*; BPGM*; PGD; PGLS; RBKS; IDH1*; ACAA1; ACAT2; JUN; FOXO1*; GCLC*; SOD2; CD80; CTLA-4; PD-1; ACE2; TNFSF10; NRG1; BIRC5; IGFBP3; CCNB2; RAB3A; SNAP25; STXBP1; Caspase 3] | 38 |
| nsp2 | [CCR8; GLUT9; ATP2A2; MYO1G; IL-2; IL-13; IL-4; UQCRB; MVD; FOXO1*; HMOX2*; FAS; ACE2; IL-18; RRAS; CTSD; CCNB2; SNAP25; STXBP1; STXBP5] | 20 |
| nsp4 | [DKK1; MT1G; CXCL11; IFI44; CXCL10; IFNλ; CCR1; CCR5; SLC6A19; TAT1; OAT1; ENO3; MYLK2; IL-13; IL-4; ENO2*; LDHA; PGD; PGLS; RBKS; UQCRB; IDH1*; ACAT2; MVD; SQLE*; JUN; HMOX2*; ERN1; PD-1; ferritin h; FAS; ACE2; CXCL2; NRG1; C5; CCL4; CTSB; CTSD; IGFBP3; STEAP3; TGFB2; CCL3L1; CXCL6; CCL8; CCL2; CCL3; SNAP25; STXBP1; Caspase 3] | 49 |
| nsp5 | [SLC6A19; ACTB; MYH13; IL-6; IL-33; PKM*; NFKB1*; JUN; FOXO1*; Ki67; CD14] | 11 |
| nsp5_c145a | [SAMD9; IFI44; CAPZB; MYO3B; PFKP; ENO2*; ALDOA; PGK1; PGD; PGLS; UQCRB; IDH1*; ACAA1; ACAT2; SOD2; ERN1; GDF11; NOG; CTSB; CTSD; SNCA; Caspase 3] | 22 |
| nsp6 | [ISG15; SLC34A2; SLC22A12; SLC6A19; EAAT3; PAT2; ASCT2; GLUT9; ABCG2; ATP2A2; TPM4; TPM3; MYO1G; OBSCN; IL-6; IL-13; IL-4; HK2*; ATP1A1*; UQCRB; IFITM3*; ACAA1; ACAT2; BCKDK; FDFT1; SQLE*; DGAT1; HMOX2*; SOD2; ERN1; ACE2; NRG1; TIMP1; AREG; CTSB; CTSD; STEAP3; SNAP25; SNCA; STXBP5] | 40 |
| nsp7 | [ISG15; IFNλ; IL-3; Megalin; SLC22A12; SLC34A1; SLC6A19; EAAT3; PAT2; ASCT2; SLC3A1; GLUT9; ABCG2; ATP2A2; ACTB; CALM3; CALM2; MYO1G; MYH15; IL-2; IL-13; IL-4; ENO2*; PKM*; LDHA; DNAJC15*; ATP1A1*; UQCRB; FDFT1; MTTP; JUN; HMOX2*; CD80; CD86; CTLA-4; Ki67; FAS; CTSD; CCL; RAB27A; RAB3A; RIMS1; RPH3A; SNCA; Tau] | 45 |
| nsp8 | [RN7SK; ISG15; PAT2; IL-12; JUN; Ki67; Tau] | 7 |
| nsp9 | [Megalin; SLC6A19; PAT2; OBSCN; MYO1A; IL-6; PGD; JUN; GCLC*; ERN1; ATF4; CXCL2; TIMP1; TGFB2; CXCL6; CCL8; CCL2] | 17 |
| orf10 | [SLC6A19; PAT2; ACTB; MYL6; CKM; IL-2; IL-6; IL-4; ENO2*; LDHA; ATP1A1*; JUN; IL-7; ACE2; TNFSF10; CTSD; CXCL1; Tau] | 35 |

| | | |
|---|---|---|
| orf3a | [CCR5; Megalin; SLC6A19; PAT2; ASCT2; ABCG2; TNNI3; TTN; OBSCN; IL-13; RBKS; FDFT1; DGAT1; ERN1; CD80; Ki67; FAS; NOG; TNFSF10; NRG1; CMTM2; STEAP3; SNAP25; STX1A; STXBP1; STXBP5; Caspase 3] | 27 |
| orf3b | [IFI44; SLC6A19; EAAT3; PAT2; ASCT2; GLUT9; ABCG2; ATP2A2; ENO3; ACTB; TPM4; TPM1; MYL6; MYH11; MYO1G; MYO7A; IL-3; GPI; LDHA; ALDOA; PGK1; PGD; ATP1A1*; UQCRB; IDH1*; CDK1*; IFITM3*; ACAA1; ACAT2; MVD; FDFT1; JUN; HMOX2*; SOD2; Ki67; SNCA] | 36 |
| orf6 | [CCR2; SNAT3; TAT1; ATP2A2; ENO3; CKM; TPM3; TMOD1; HK2*; ALDOA; PGK1; PGD; ATP1A1*; UQCRB; IDH1*; ACAT2; JUN; SOD2; CD80; PD-1; FAS; CD68; NRG1; CTSD; RRM2; TGFB2; UNC13A; RAB3A] | 28 |
| orf7a | [CCR2; PAT2; ATP2A2; MYLK2; IL-13; PFKP; PKM*; ACAT2; BCKDK; JUN; DNAJB9; CD80; Ki67; FAS; ACE2; CD68; NRG1; IL-18; STEAP3; Tau] | 20 |
| orf8 | [DKK1; IFI44L; IFI44; IL-3; IL-5; SLC6A19; ASCT2; GLUT9; MYH15; MYO1F; TNF-α; IL-1β; IL-2; IL-6; IL-12; IL-13; IL-33; IL-4; IL-10; IL-17; HK2*; MTTP; SOD2; ERN1; CD80; CRP; ACE2; MRC1 cd206; GDF11; NOG; NRG1; TIMP1; CTSD; IGFBP3; CCNB2; TGFB2; CCL7; SNCA] | 38 |
| orf9b | [IFI44L; RSAD2; IL-3; SLC22A12; SLC34A1; SLC6A19; PAT2; ABCG2; ENO3; TNNC1; CKM; TNNI3; CAPZB; IKKβ; IL-6; IL-33; IL-4; HK2*; PKM*; PGD; ACAT2; MVD; JUN; GCLC*; ERN1; CD80; CTLA-4; ACE2; TNFSF10; CTSB; DDIT4; IGFBP3; CCNB2; TGFB2; RIMS1; SNAP25; STXBP5; Tau] | 38 |
| orf9c | [MX1; DKK1; IFNλ; CCR1; CCR8; SLC22A12; EAAT3; SNAT3; TAT1; OAT1; OAT4; GLUT9; ATP2A2; ACTB; MYH6; MYH15; IL-2; IL-13; IL-4; IL-10; IL-17; IL-22; SQLE*; DGAT1; HMOX2*'; ERN1; DNAJB9; CD80; IL-7; PD-1; FAS; ACE2; NOG; NRG1; C5; CCL4; CMTM2; CCL3; CXCL9; RAB3A; STXBP5] | 41 |
